# Explaining the primate extinction crisis: predictors of extinction risk and active threats

**DOI:** 10.1101/2022.12.08.519299

**Authors:** Maria J.A. Creighton, Charles L. Nunn

## Abstract

Explaining why some species are disproportionately impacted by the extinction crisis is of critical importance for conservation biology as a science and for proactively protecting species that are likely to become threatened in the future. Using the most current data on threat status, population trends, and threat types for 446 primate species, we advance previous research on the determinants of extinction risk by including a wider array of phenotypic traits as predictors, filling gaps in these trait data using multiple imputation, and investigating the mechanisms that connect organismal traits to extinction risk. Our Bayesian phylogenetically controlled analyses reveal that insular species exhibit higher threat status, while those that are more omnivorous and live in larger groups have lower threat status. The same traits are not linked to risk when repeating our analyses with older IUCN data, which may suggest that the traits influencing species risk are changing as anthropogenic effects continue to transform natural landscapes. We also show that non-insular, larger-bodied, and arboreal species are more susceptible to key threats responsible for primate population declines. Collectively, these results provide new insights to the determinants of primate extinction and identify the mechanisms (i.e., threats) that link traits to extinction risk.

## INTRODUCTION

Anthropogenic activity is causing species to disappear at an alarming rate. However, not all species are affected equally. Explaining why some species are more susceptible to extinction than others has become a major goal of conservation biologists as these contributions help to both explain current extinction patterns and allow for proactive protection of species possessing traits that could increase their probability of becoming imperiled. Previous studies have shown that phenotypic traits affect a species’ susceptibility to extinction (Chichorro et al., 2019). Physical traits such as large body size and life history traits such as long generation lengths have been associated with increased risk of extinction in some clades (Bennett & Owens, 1997; Purvis et al., 2000; Cardillo & Bromham, 2001; Cardillo et al., 2005; Fritz, et al., 2009; Lee & Jetz, 2011; Matthews et al., 2011; Ripple et al., 2017; Nolte et al., 2019; Chichorro et al., 2022a; Chichorro et al., 2022b). These findings match expectations that lower population densities and increased hunting pressures put larger species disproportionately at risk and expectations that species with longer life histories have less time to adapt to environmental changes (Purvis et al., 2000; Cardillo & Bromham, 2001; Cardillo et al., 2005; Cardillo, 2021). Behavioural traits have also been linked to increased extinction risk, including small group size and reduced innovativeness (Davidson et al., 2009; 2012; Ducatez et al., 2020): large groups are expected to benefit from reduced predation and enhanced foraging while less innovative species are less well-equipped to solve novel environmental challenges.

While much effort has been put toward identifying the traits that covary with extinction risk, important knowledge gaps have limited the effectiveness of these analyses. First, only a handful of studies have incorporated a broad range of traits in a single analysis. Chichorro et al. (2019) reviewed studies investigating the correlates of extinction risk and found significant variability in the traits that were investigated (or controlled for). In addition, some traits have only recently been linked to extinction risk, such as behavioural flexibility (Ducatez et al., 2020), and thus have not been widely investigated across clades.

Second, the relationship between the actual anthropogenic drivers of environmental change that are responsible for extinction and species traits are understudied in many clades (e.g., in primates; Estrada et al., 2017), limiting the impact of these comparative studies in applied conservation (Cardillo & Meijaard, 2012). Identifying which threats are most impactful to species with different trait types would enable actionable conservation steps (e.g., mitigating key threats in susceptible species’ ranges). Despite the possible benefit of considering specific threats, previous research has mostly focused on connecting species’ traits and threat status. Notably, some studies focused primarily on predictors of threat status have attempted to incorporate information about anthropogenic threats into their analyses (e.g., Purvis et al., 2005; Cardillo et al., 2008; González-Suárez et al., 2013; Murray et al., 2014; Di Marco et al., 2015; Ruland & Jeschke, 2017; Atwood et al., 2020; Chichorro et al., 2022a) while Richards et al. (2021) recently explicitly assessed predictors of anthropogenetic threats to seabirds.

Lastly, we lack information on relevant traits for many species, resulting in incomplete data. The species for which we lack data may be systematically biased towards those that are more difficult to study, such as arboreal or nocturnal species. In addition to reducing statistical power, removing these species from analyses has potential to bias observed relationships between variables (Nakagawa & Freckleton, 2008) and can result in a loss of real information when some traits included in an analysis have better data coverage than others. In recent years, improved methods for imputing missing data have become available, creating opportunities to reduce the number of missing data points in analyses of extinction risk (e.g., see Richards et al., 2021; Chichorro et al., 2022b).

Primates have been especially important in studies assessing predictors of extinction risk (Purvis et al., 2000; Purvis et al., 2005; Matthews et al., 2011; Machado et al., 2022). Primates are crucial components of tropical biodiversity, core players in the function of ecosystems, and central to many cultures and religions (Estrada et al., 2017). It is thus an urgent goal to determine which biological and behavioural traits contribute to primate extinction vulnerability and how these traits interact with anthropogenic impacts to contribute to population declines.

Primates are also one of the most threatened animal clades, with ∼65% of species at risk of extinction (IUCN, 2021), yet the last comprehensive assessment of the major determinants of primate extinction risk was published over 20 years ago (Purvis et al., 2000). The number of recognized primate species has changed dramatically since earlier studies, having more than doubled from 180 to over 500 in the past few decades (Rylands & Mittermeier, 2014; Creighton et al., 2022). As a result of these taxonomic changes and limitations of older phylogenies, older studies focused on a relatively small number of currently recognized primate species. More speciose and up-to-date phylogenies have recently become available (Upham et al., 2019), coupled with a greater quantity and quality of trait data for many primate species. These contributions create an opportunity for the inclusion of more described primate species in comparative analyses, bringing us closer to capturing the true scope of primate diversity.

Here, we analyze the biological and behavioural determinants of primate extinction risk using a phylogenetic comparative approach. We investigate the relationship between multiple phenotypic traits and two measures of extinction risk reported by the International Union for Conservation of Nature (IUCN): threat status and population trend. We then assess how these same traits covary with vulnerability to the major threats facing primate species. This research addresses the gaps above by including multiple traits in the analysis and using imputation approaches based on phylogeny and phenotypic traits to fill in data for species with missing trait values. In addition, by investigating population trends and specific threats, we improve understanding of the connections between specific traits and the abundance of primates.

We focus on 10 key traits with proposed links to extinction risk (Table 1).

**Table 1:**
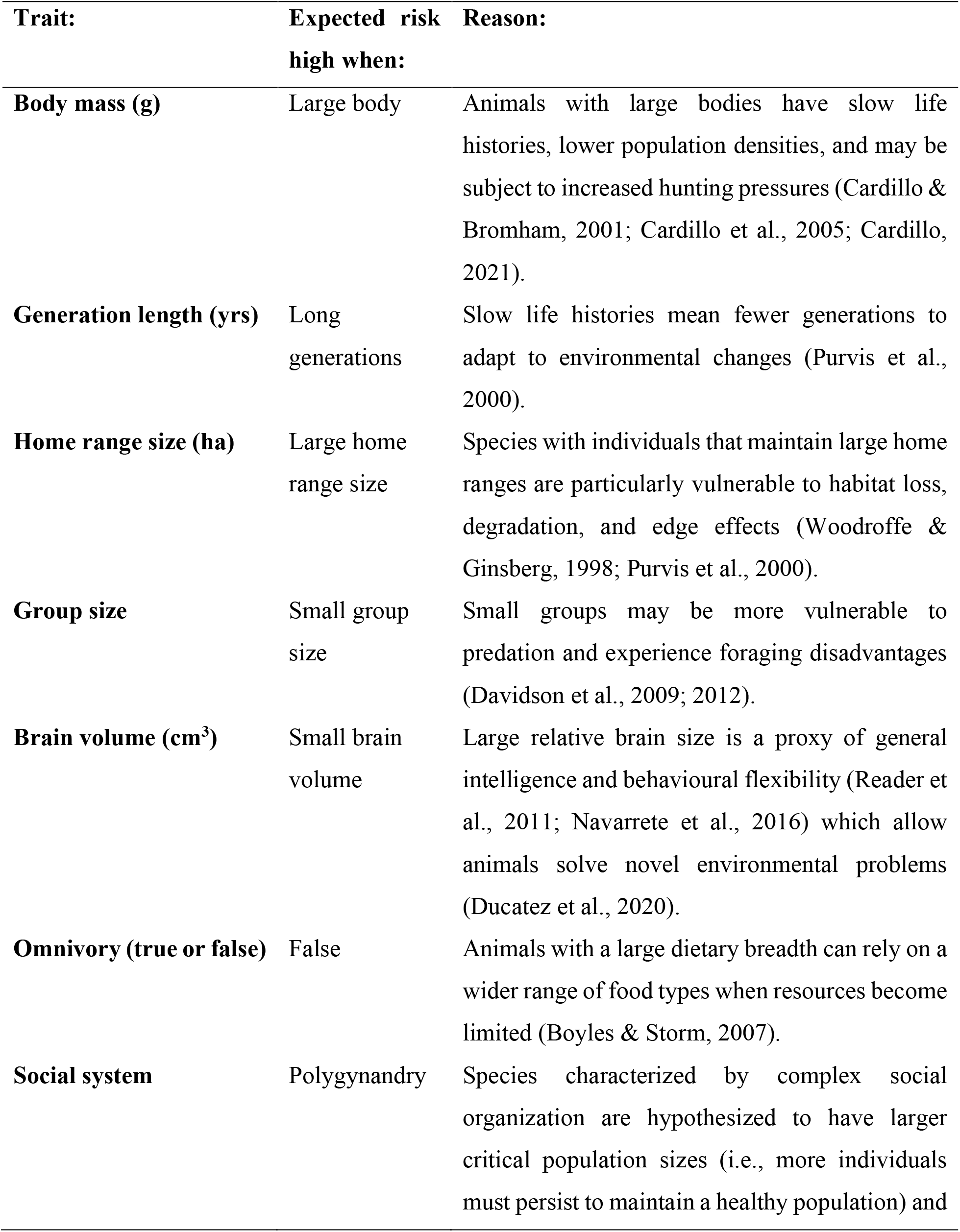
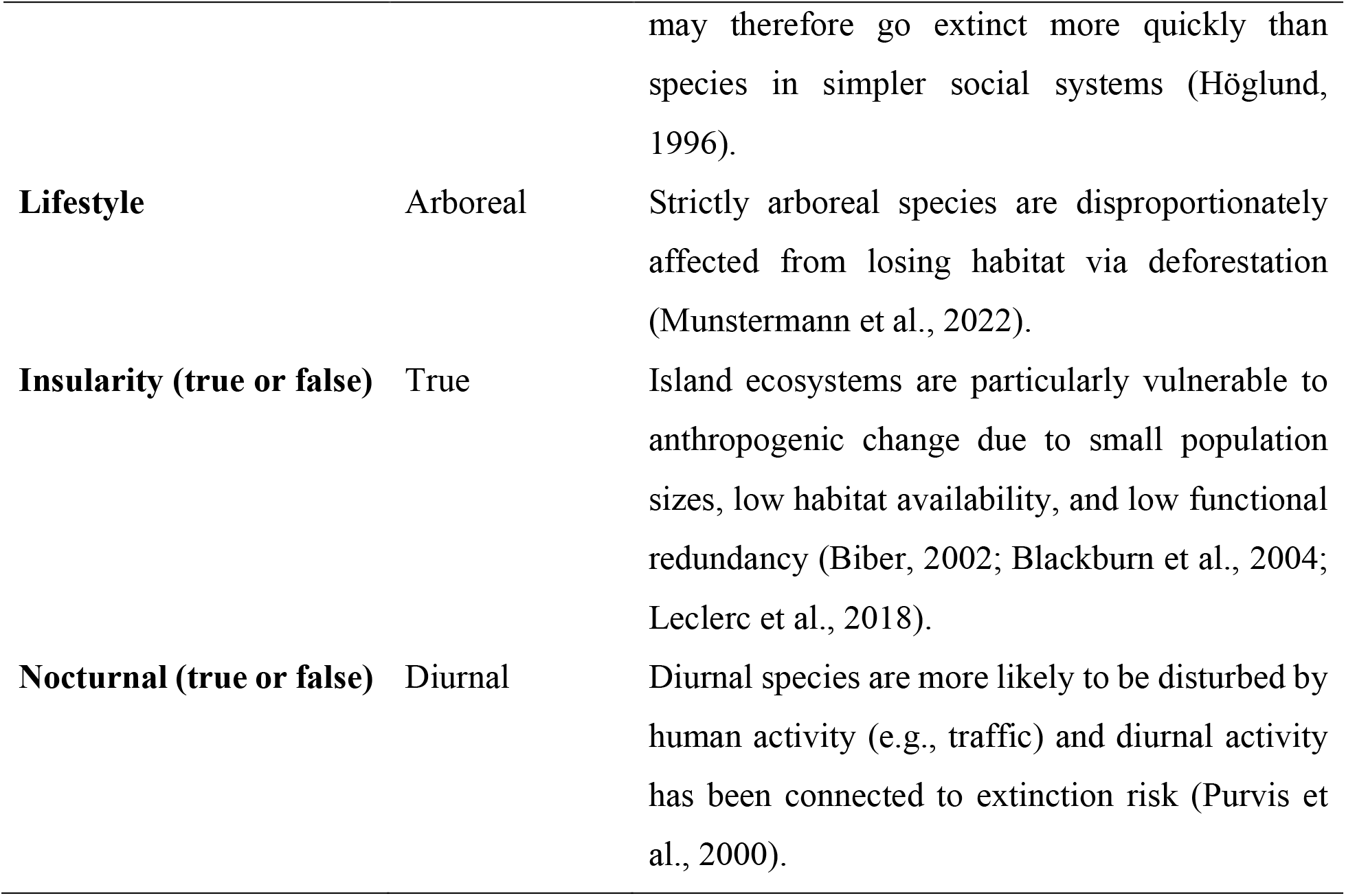
The predicted direction of effect of biological and behavioural traits on extinction risk.

| **Trait:** | **Expected risk high when:** | **Reason:** |
| --- | --- | --- |
| **Body mass (g)** | Large body | Animals with large bodies have slow life histories, lower population densities, and may be subject to increased hunting pressures (Cardillo & Bromham, 2001; Cardillo et al., 2005; Cardillo, 2021). |
| **Generation length (yrs)** | Long generations | Slow life histories mean fewer generations to adapt to environmental changes (Purvis et al., 2000). |
| **Home range size (ha)** | Large home range size | Species with individuals that maintain large home ranges are particularly vulnerable to habitat loss, degradation, and edge effects (Woodroffe & Ginsberg, 1998; Purvis et al., 2000). |
| **Group size** | Small group size | Small groups may be more vulnerable to predation and experience foraging disadvantages (Davidson et al., 2009; 2012). |
| **Brain volume (cm<sup>3</sup>)** | Small brain volume | Large relative brain size is a proxy of general intelligence and behavioural flexibility (Reader et al., 2011; Navarrete et al., 2016) which allow animals solve novel environmental problems (Ducatez et al., 2020). |
| **Omnivory (true or false)** | False | Animals with a large dietary breadth can rely on a wider range of food types when resources become limited (Boyles & Storm, 2007). |
| **Social system** | Polygynandry | Species characterized by complex social organization are hypothesized to have larger critical population sizes (i.e., more individuals must persist to maintain a healthy population) and |
|  |  | may therefore go extinct more quickly than species in simpler social systems (Höglund, 1996). |
| **Lifestyle** | Arboreal | Strictly arboreal species are disproportionately affected from losing habitat via deforestation (Munstermann et al., 2022). |
| **Insularity (true or false)** | True | Island ecosystems are particularly vulnerable to anthropogenic change due to small population sizes, low habitat availability, and low functional redundancy (Biber, 2002; Blackburn et al., 2004; Leclerc et al., 2018). |
| **Nocturnal (true or false)** | Diurnal | Diurnal species are more likely to be disturbed by human activity (e.g., traffic) and diurnal activity has been connected to extinction risk (Purvis et al., 2000). |

## METHODS

### DATA

We collected information on threat status (least concern = LC, near threatened = NT, vulnerable = VU, endangered = EN, critically endangered = CR, data deficient = DD, and not evaluated = NE) and population trend (increasing = I, stable = S, decreasing = D, and unknown = U), from the IUCN (2021) for 446 primate species present in the ultrametric primate phylogeny published by Upham et al. (2019) (Figure 1). We also collected a list of active threat types affecting each species in the IUCN (2021) as defined by the Salafsky et al. (2008) threat classification system: 1 = residential and commercial development, 2 = agriculture and aquaculture, 3 = energy production and mining, 4 = transportation and service corridors, 5 = biological resource use, 6 = human intrusions and disturbance, 7 = natural system modifications, 8 = invasive and other problematic species and genes, 9 = pollution, 10 = geological events, and 11 = climate change and severe weather.

**Figure 1.**
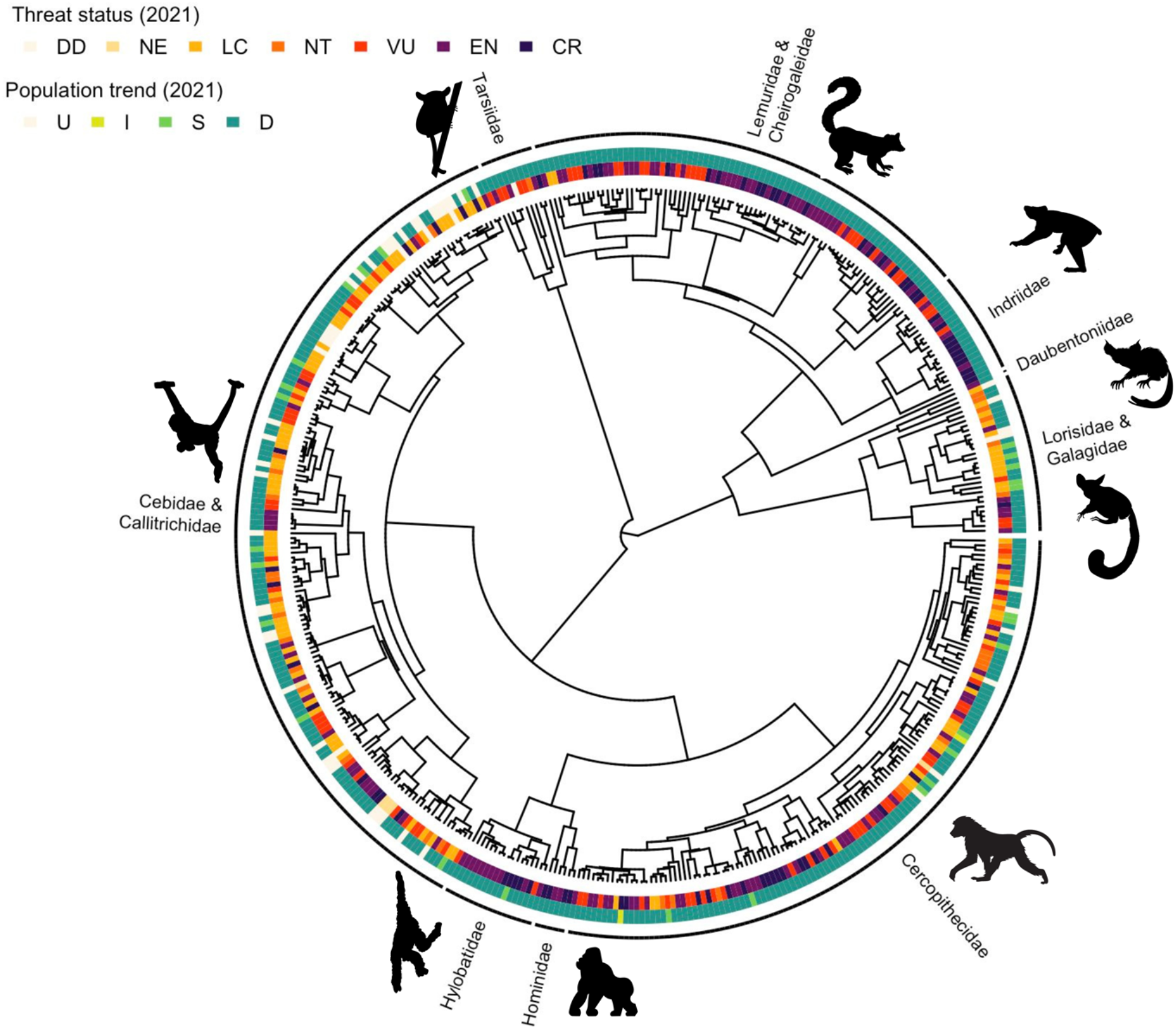
Phylogenetic distribution of threat status and population trends (IUCN, 2021) for 446 primate species in the Upham et al. (2019) phylogeny. Images of representative species are presented next to family labels. Codes for threat status: data deficient = DD, not evaluated = NE, least concern = LC, near threatened = NT, vulnerable = VU, endangered = EN, and critically endangered = CR. Codes for population trend: unknown = U, increasing = I, stable = S, and decreasing = D (IUCN, 2021).

For each of the 446 species in our dataset, we recorded data on 10 different biological and behavioural traits that have been proposed to be associated with extinction risk from various sources: body mass (g) (Galán-Acedo et al., 2019), generation length (yrs) (IUCN, 2021), home range size (ha) (Galán-Acedo et al., 2019), group size (Rowe & Myers, 2011), brain volume (Powell et al., 2017), omnivory (true or false), social system (solitary, pair-living, harem polygyny and polygynandry) (DeCasien et al., 2017; Rowe & Myers, 2011 and other sources), lifestyle (arboreal, terrestrial, or both) (Rowe & Myers, 2011 and other sources), insular (true or false) (inferred from range data available in Rowe & Myers, 2011; IUCN, 2021), and nocturnal (true or false) (Estrada et al., 2017; IUCN, 2021). The full list of references for trait values is available in the Supplementary Data. Table 1 summarizes how we expected each trait to be associated with primate extinction risk. Previous studies have included geographic range size as a covariate in similar analyses (e.g., Purvis et al., 2000; Machado et al., 2022). However, a species’ geographic range size is one of the main criteria used by the IUCN to assign threat status: species with small population sizes that have small or restricted geographic ranges are considered to be more imperiled (IUCN, 2021) (i.e., threatened species have small geographic ranges by definition). Because we were interested in how biological and behavioural traits contribute to extinction risk, including effects on what geographic ranges they can occupy, we did not include geographic range size in our analysis. Notably, by including insularity in our analysis we controlled for the fact that species on small islands may not be able to maintain geographic ranges large enough to be considered healthy by the IUCN due to geographic barriers.

Following Powell et al. (2017), for sexually dimorphic clades (size difference > 10%) only brain volume and body mass data from adult females were used in analysis. For all other species, averages for all adults measured in the original source were used. Species found exclusively on Madagascar, Borneo, or Sumatra were not scored as insular since these islands are large enough to support large geographic ranges comparable to many mainland species. Further details on operational definitions and trait coding are provided in the Supplementary Materials, along with a correlation matrix of all traits and response variables (Figure S1), and a comparison of trait data from different sources (Figures S2, S3, and S4).

### ANALYSIS

#### Trait imputation

The availability of data varied across the species in our dataset. Percentages of species with missing trait data were: body mass (6%), omnivory (17%), generation length (23%), home range size (25%), group size (39%), and brain volume (46%). Restricting the analysis to only species with observed data on all traits reduced our sample size of species by over half (e.g., to from n=430 to n=151 in our analysis of threat status). We thus opted to use multiple imputation for these six traits to avoid losing species from our analysis where one or more traits had missing observations. The other four traits in our analysis (lifestyle, nocturnality, insularity, and social system) had full data coverage and no imputation procedure was necessary for these traits.

Multiple imputation was accomplished using 100 randomly sampled trees available from Upham et al. (2019). From each tree, we first generated a variance-covariance matrix which we then dissolved into 445 eigenvectors using the ‘PVRdecomp’ function from the R package PVR (Santos et al., 2018). To impute body mass (the trait with the best data coverage requiring imputation), we built a linear model where body mass was the response and phylogenetic eigenvectors were predictors. Using forward-backward model selection, we then determined which phylogenetic eigenvectors had the best support for inclusion in models predicting body mass based on Akaike information criterion (AICc) scores using the ‘stepAIC’ function from MASS (Ripley et al., 2013). We chose how many eigenvectors to include in model selection for the imputation based on model performance in cross validation (Table S1). We repeated this process for all traits with missing data. The imputation of traits was ordered so that imputed information could be used to inform subsequent model fits along with phylogenetic information (e.g., once body mass was imputed it was used to inform model fits for the imputation of other traits). Body mass (including both sexes) and female body mass were imputed separately, and female body mass was used thereafter as body mass for sexually dimorphic clades. The top model for each trait was used to impute values for each species with missing data using the ‘predict’ function from the car package (Fox et al., 2012). To propagate error, we then used the fits and standard deviations associated with predicted values to take a randomly sampled trait value for each species from the normal or binomial distribution (depending on whether traits were continuous or binary).

The imputation of traits was repeated once for each tree, resulting in 100 imputed datasets. We performed a leave-one-out cross-validation of each imputation, where we removed observed datapoints and used our imputation method to predict their values. When comparing these predictions to the observed datapoints performance proved to be good in all cases (predictive accuracy > 0.8 for continuous variables and area under the ROC curve > 0.8 for binary variables; see Table S1).

Different methods of trait imputation can lead to different trait estimates, which may affect conclusions in downstream analyses. To assess the consistency of our results using different methods, we ran a second imputation procedure with the ‘phylopars’ function from Rphylopars (Goolsby et al., 2017). Rphylopars uses a maximum likelihood approach with a phylogeny and sparse trait matrix to impute missing data (Goolsby et al., 2017; Johnson et al., 2021). Rphylopars provides an advantage over the previously described imputation approach by incorporating the best supported model of trait evolution (in our case the Ornstein–Uhlenbeck model) to impute missing traits values (Goolsby et al., 2017), whereas our other imputation method did not subscribe to a particular model of trait evolution. However, Rphylopars also has some limitations including a lack of customizability (e.g., optimizing model inputs based on cross-validation performance) and can still lead to biased trait estimates particularly for traits with a large amount of missing data (Johnson et al., 2021). Given the advantages and disadvantages of each approach we decided run our main analyses with 100 sets of imputed data from both imputation methods to inform our conclusions about which traits are associated with primate extinction risk.

#### Modelling threat status, population trends, and threat types

We ran multiple models to test predictors of three types of outcomes for primate populations. First, we tested which biological and behavioural traits are associated with the threat status of a species. Second, we tested the effects of species’ traits on population trends. Third, we tested which traits were associated with species susceptibility to the most prevalent threats that primate species face.

To determine which biological and behavioural traits are associated with threat status, we ran two phylogenetic generalized linear mixed models using Bayesian approximation as implemented in the MCMCglmm R package (Hadfield, 2010). The first model had an ordinal error structure, and the response variable was an ordinal measure of threat status scored as follows: LC=0, NT=1, VU=2, EN=3, and CR=4 (Butchart et al., 2007). We ran the second model with a threshold error structure; here the response variable was threat status scored as a binary outcome where species were scored as either being threatened (VU, EN, or CR) (scored as 1) or not threatened (NT or LC) (scored as 0).

To test the effects of traits on population trends we ran a phylogenetic generalized linear mixed model with a threshold error structure (Hadfield, 2010). In this model, each species was assigned a binary outcome of either declining (scored as 1) or not declining (i.e., stable or increasing) (scored as 0).

Finally, to determine which biological and behavioural traits were associated with species’ susceptibility to the most important threats that primate species face, we ran five phylogenetic generalized linear mixed models with threshold error structures (Hadfield, 2010), one for each of the top five threats to primates identified by the IUCN (2021). These top five threats identified for primate species were: 1 = residential and commercial development (35% of species), 2 = agriculture and aquaculture (80%), 3 = energy production and mining (22%), 5 = biological resource use (82%), and 7 = natural system modifications (23%). Here, each of our five models had a binary outcome of 1 (indicating that a species was affected by a particular threat) or 0 (indicating that a species was not affected by the threat).

Each model described above included all 10 traits of interest as fixed effects (i.e., traits were included together as predictors in the same model) and each model was run on 100 imputed datasets and phylogenies to account for uncertainty in phylogeny and trait estimates (Nakagawa & De Villemereuil, 2019). Models ran a total of 550,001 iterations, with a thinning interval of 500 and a burn-in of 50,000 to ensure convergence had occurred, which we assessed using trace plots (Hadfield, 2010). Species were dropped from analyses when the true value of a response variable was unknown by the IUCN (2021) (e.g., if the threat status was DD or NE). Continuous variables were ln-transformed, centered with respect to the mean, and scaled by 2 standard deviations in all models to make their effect sizes comparable to those reported for binary variables (Gelman, 2008). We used a weakly informative gelman prior for fixed effects and fixed the residual variance (R) to 1 (Hadfield, 2010). For the phylogenetic component (G), we used parameter expanded priors where V = 1, nu = 1, alpha.mu = 0, and alpha.V = 100 (Hadfield, 2018). These analyses were repeated using 100 imputed datasets from both our model selection approach and Rphylopars.

We also tested the hypothesis that some traits previously shown to be associated with primate extinction risk are losing signal as more species become imperiled, for example, if anthropogenic threats are becoming so overwhelming that all species are beginning to suffer regardless of their attributes. This analysis involved repeating our analyses of threat status using an older IUCN threat status data and species list (obtained from Harcourt & Parks, 2003).

To interpret the output from our Bayesian analyses, we provide (i) the distribution of posterior means for tests from all 100 imputed datasets in graphical form (Figures 2 and 3), (ii) the 89% credible intervals (per McElreath, 2018) from the full posterior distribution of estimates in graphical form (Figures 2 and 3), and (iii) the percentage of iterations from each set of 100 models that were consistently positive or negative (Tables S2 to S20). We focused on results that were most supported based on these outcomes and supported consistently using data from both imputation procedures. For the purposes of providing an estimate of the magnitude of an effect in the main text, posterior means were pooled across datasets using Rubin’s rules (Nakagawa & De Villemereuil, 2019) (hereafter, “pooled posterior mean”). Results reported in the main text are from our model selection imputation procedure and were qualitatively consistent using Rphylopars unless otherwise stated.

**Figure 2.**
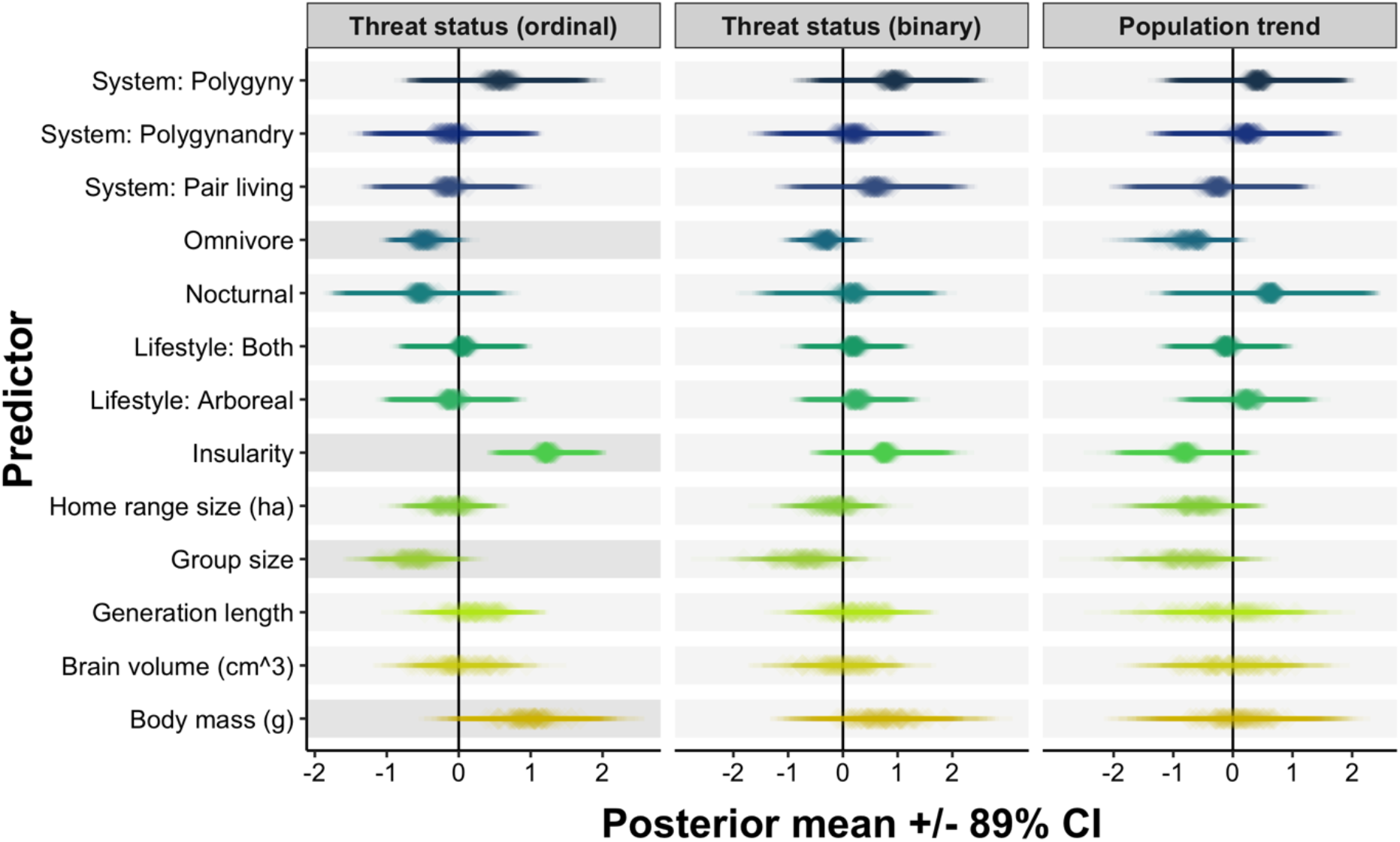
Outcomes from three sets of models testing the relationship between traits and: threat status scored as an ordinal variable (first panel), threat status scored as a binary variable (second panel), and population trend scored as a binary variable (third panel). Each cell contains 100 posterior means (plotted as translucent diamonds) with their associated 89% credible intervals (plotted as translucent horizontal lines) obtained from 100 MCMCglmm models run with 100 randomly sampled phylogenies across 100 trait datasets with missing datapoints obtained through multiple imputation. Darker shading behind effects interpreted in text. Continuous variables were ln-transformed, centered with respect to the mean, and scaled by 2 standard deviations.

**Figure 3.**
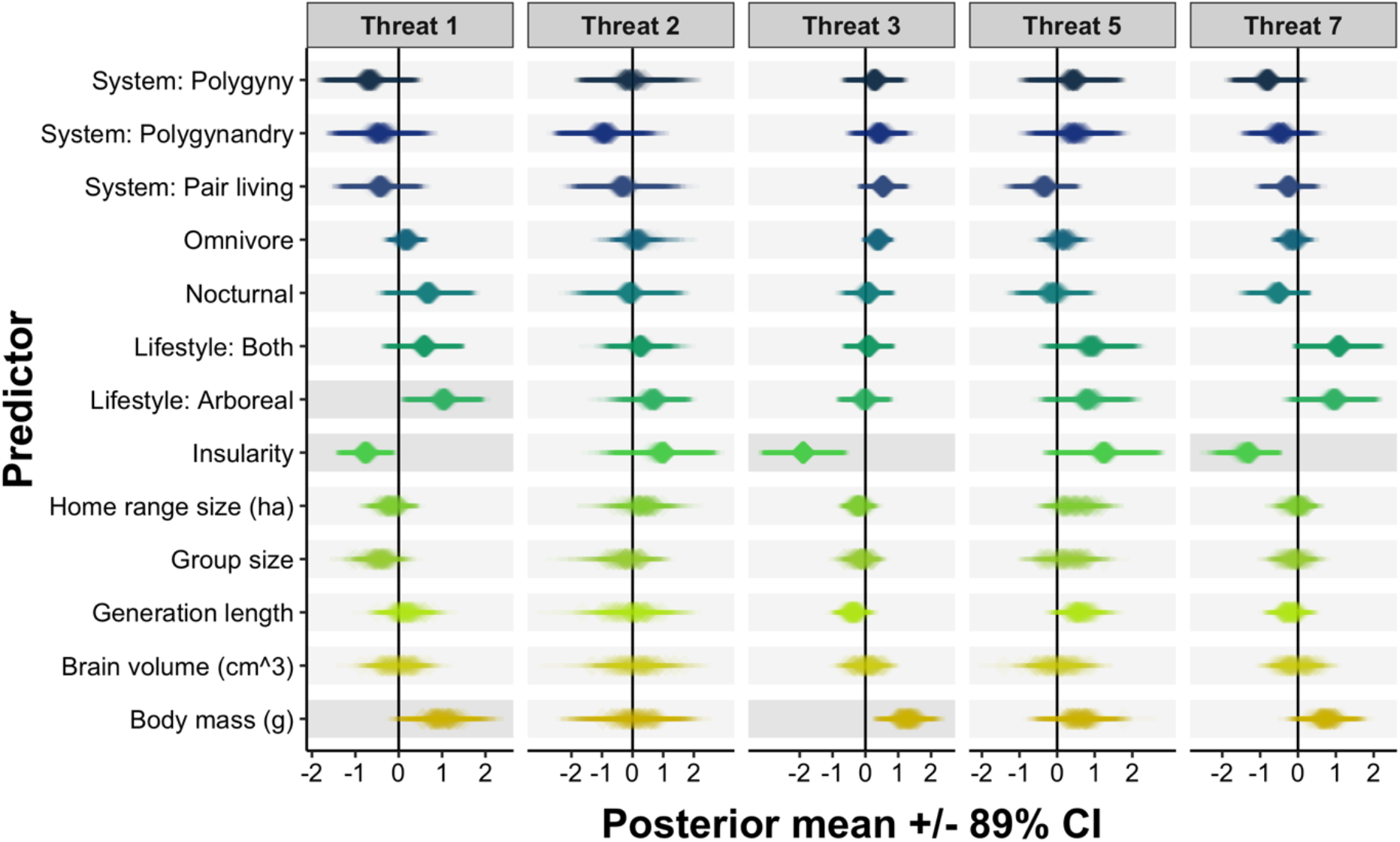
Outcomes from five sets of models testing the relationship between traits and species susceptibility to: threat 1 = residential and commercial development (first panel), threat 2 = agriculture and aquaculture (second panel), threat 3 = energy production and mining (third panel), threat 5 = biological resource use (fourth panel), and threat 7 = natural system modifications (fifth panel). Each cell contains 100 posterior means (plotted as translucent diamonds) with their associated 89% credible intervals (plotted as translucent horizontal lines) obtained from 100 MCMCglmm models run with 100 randomly sampled phylogenies across 100 trait datasets with missing datapoints obtained through multiple imputation. Darker shading behind effects interpreted in text. Continuous variables were ln-transformed, centered with respect to the mean, and scaled by 2 standard deviations.

## RESULTS

### Predictors of threat status and population trends

Our dataset included 430 primate species with known threat statuses. When scored as an ordinal outcome (LC=0, NT=1, VU=2, EN=3, and CR=4), primate threat status was positively associated with insularity (pooled posterior mean = 1.214; 100% of 100,100 posterior estimates > 0) (Figure 2; Tables S2). Ordinal threat status was negatively associated with omnivory (pooled posterior mean = -0.474; 95% estimates < 0) and group size (pooled posterior mean = -0.561; 94% estimates < 0) (Figure 2; Table S2). Body mass, which is often considered a main predictor of extinction risk in many clades, only weakly predicted threat status after controlling for other covariates (pooled posterior mean = 0.980; 93% estimates > 0) and this weak support disappeared when using imputed data from Rphylopars (pooled posterior mean = 0.517; 74% estimates > 0) (Tables S8). In our analysis using older IUCN data, we found that only insularity (pooled posterior mean = 2.255; 100% estimates > 0) and home range size (pooled posterior mean = 1.498; 100% estimates > 0) were associated with threat status (ordinal) in the predicted direction (Table S3).

Threat status was not strongly associated with any biological or behavioural traits when scored as a binary response (Figure 2; Table S4). We also ran three separate models with body mass, generation length, and brain volume as sole predictors to determine if correlations among these variables (see Figure S1) contributed to the lack of strong associations. However, these additional analyses consistently showed no strong effect of any traits (Tables S5). In our analysis using older IUCN data, we found that insularity (pooled posterior mean = 2.556; 100% estimates > 0) and home range size (pooled posterior mean = 1.078; 99% estimates > 0) were associated with threat status (binary) in the predicted direction (Table S6).

Population trend was not consistently associated with any biological or behavioural traits across 401 species with known population trends (Figure 2; Table S7).

### Predictors of specific threat types

Our analyses of threat types included 404 species with known threats. Insularity was negatively associated with threat 1 = residential and commercial development (pooled posterior mean = -0.752; 98% estimates < 0; Table S11), threat 3 = energy production and mining (pooled posterior mean = -1.807; 100% estimates < 0; Table S13), and threat 7 = natural system modifications (pooled posterior mean = -1.312; 100% estimates < 0; Table S15) (Figure 3). Species living a strictly arboreal lifestyle were more likely to be affected by threat 1 = residential and commercial development than strictly terrestrial species (i.e., the baseline) (pooled posterior mean = 1.045; 98% estimates > 0; Table S11) (Figure 3). Species with larger body masses were more likely to be affected by threat 3 = energy production and mining (pooled posterior mean = 1.232; 99% estimates > 0; Table S13) (Figure 3). Large bodied species were also more likely to be affected by threat 1 = residential and commercial development (pooled posterior mean = 1.004; 96% estimates > 0; Table S11), but like our analysis of ordinal threat status, this effect was no longer compelling using imputed data from Rphylopars (pooled posterior mean = 0.741; 83% estimates > 0; Table S16) which instead supported a negative relationship between threat 1 and group size (pooled posterior mean = -0.720; 97% estimates < 0; Table S16).

Some weaker trait associations were also detected (e.g., arboreality or partial arboreality being positively associated with threat 7; Table S15). However, given the number of models where credible intervals around these estimates overlapped with zero, we chose to interpret associations with stronger support.

## DISCUSSION

We investigated the correlates of extinction risk and threat susceptibility in primates using phylogenetic comparative methods to analyze the most complete and up-to-date set of trait data and IUCN data. One novelty of our approach was the use of phylogenetic and trait-based imputation of missing data. In our analysis using threat status as ordinal outcome – with five ranked categories from least concern to critically endangered – higher threat status was associated with insularity, omnivory, and small group size, consistent with our predictions for these traits. Therefore, primate species that are most imperiled, and thus score highest in ordinal threat status, do tend to be those with biological and behavioural predispositions to extinction. When looking at specific threats, we found that larger-bodied and arboreal species are more vulnerable to key threats, while insular species are less vulnerable to these threats.

Although we did find that some traits were powerful predictors of threat status when scored according to ranked categories, the traits we investigated were not strong predictors of binary extinction risk outcomes (i.e., threat status scored as threatened versus non-threatened and population trend scored as declining versus not declining). This is contrary to findings in other taxonomic groups, such as birds, where multiple traits have been linked to binary measures of extinction risk (Lee & Jetz, 2011). Notably, the relative number of threatened and declining species is much higher for primates than for birds. For instance, 93% of primate species with known population trends are in decline while ∼50% of birds are in decline (Lee & Jetz, 2011). This may explain why no predictors emerge for binary outcomes in primates: most species are threatened or becoming threatened, and traits are mostly deterministic of severity.

Some traits that predict threat status in other clades did not emerge as powerful predictors for primates, even in our analysis of ordinal threat status. For example, in a recent study of birds, Ducatez et al. (2020) provided evidence that innovativeness – a known measure of behavioural flexibility associated with general intelligence (Reader & Laland, 2002; Reader et al., 2011) – buffers against extinction. However, we did not find an effect of brain size, another known measure of behavioural flexibility (Reader et al., 2011; Creighton et al., 2021) and a correlate of innovativeness, in our analyses. The caveats associated with each of these measures are discussed in Creighton et al. (2021), however, we suggest that these differences in results could be because the links between extinction risk and flexibility are complex for a clade like primates where innovations are frequent and human conflict is common. Certain behavioural innovations may reduce conflict with humans and increase resilience to habitat degradation (e.g., novel approaches for accessing food; Beever et al., 2017). However, other innovative behaviours can increase human-wildlife conflict. In many primate species, crop-raiding and garbage eating are innovative behaviours that have become common practice in the context of anthropogenic change (e.g., chimpanzees and baboons; Maples et al., 1976; Hahn et al., 2003; Hockings et al., 2009). These behaviours bring animals in direct conflict with humans and, in some cases, attract them to lower quality habitats (McLennan et al., 2017). Future contributions should further address the paradox of how flexibility both helps and hinders species’ persistence.

When repeating our analyses of extinction risk with a 1999 IUCN dataset, we found that insularity and home range size shared a positive relationship with binary and ordinal threat status, but other traits were not powerful predictors. This pattern of results using newer versus older data indicates that some traits (i.e., home range size) have become less powerful predictors of extinction since 1999. Meanwhile, some traits identified in our analysis of ordinal 2021 threat status (i.e., body mass, group size, and omnivory) do not emerge in analyses with older data indicating these traits may be beginning to have a larger signal over time. However, it is also possible that the larger number of species in our 2021 dataset (a consequence of taxonomic reevaluations in many clades; Creighton et al., 2022) and general improvements in the thoroughness and accuracy of IUCN assessments since 1999 provided the statistical power to detect the effects of these traits.

In analyses assessing predictors of direct threats, we found that strictly arboreal species were more likely to be threatened with residential and commercial development – a major driver of deforestation in many regions. We also found that insular species were less likely to be vulnerable to multiple threats (residential and commercial development, energy production and mining, and natural system modifications) despite being more likely to be highly threatened. This indicates that the high threat statuses of insular species may not be driven by anthropogenic activity. Instead, their small geographic ranges enforced by geographic barriers could simply make it impossible to maintain healthy population sizes, despite not being subject to major threats.

While our results were largely consistent using two methods of imputation, we identified a few discrepancies where support for trait associations changed meaningfully depending on the imputation procedure: one method better supported an effect of body mass on ordinal threat status and threat type 1, while the other better supported an effect of group size on threat 1. One of the main benefits of imputation in multivariable analyses is the ability to preserve real data for well represented traits when another trait has poor coverage. Given that all methods of imputation are imperfect, using multiple methods of imputation with different associated biases may help reveal cases where the imputation is influential in driving an observed relationship between two variables and may help interpretation of findings, especially when credible intervals are wide or close to overlapping with zero.

One limitation of our analysis was that body mass, generation length, and brain volume were all highly correlated in our dataset (correlation coefficients range between 0.6 and 0.9). These traits each have independent predicted associations with extinction risk in conflicting directions, meaning that reducing these variables to fewer terms (e.g., via principal component analysis) would reduce our ability to draw conclusions about their independent effects. We instead included these predictors in the same models to identify how they independently contributed to extinction risk and threat vulnerability (Freckleton, 2002). However, this creates the possibility of collinearity in model estimates for these variables as suggested by the increased uncertainty (i.e., wide credible intervals) around estimates for these three traits. Notably, when we did not detect an effect of these traits we tested them as predictors in separate models and results suggested that associations among these traits were not strong enough to affect our conclusions. We were also limited in that, like many previous studies, response variables in our analysis come from IUCN assessments. While the IUCN maintains the largest global dataset on species extinction risk and threats useful for comparative analyses, these measures are vulnerable to errors in empirical data and in models used to estimate population declines and extinction risk (Rueda-Cediel et al., 2018). As a result, there is likely to be uncaptured uncertainty associated with the measures of extinction risk used in our analyses.

Cardillo & Meijaard (2012) identified the limitations of comparative studies of extinction risk when it comes to conservation action, including the difficulty of translating results to policy and on the ground conservation activities. We therefore offer some connections between our findings and real-world conservation questions. First, understanding the biological and behavioural predictors of threat susceptibility in broader range of taxa could help organizations like the IUCN to identify which threats pose the most imminent harm to species with shared characteristics. When it comes to conservation action, this information can be used to identify which populations are most likely to become threatened by anthropogenic activity in the near future based on a combination of imminent threats, traits, and species’ distributions. For instance, overlaying geographical information about the expansion of threats described above on primate species’ distributions would allow us to forecast which populations are most likely to be impacted based on their traits (e.g., arboreal species living in proximity to residential developments areas may be at particularly high risk). This approach may help identify where to allocate limited funds for conservation and surveillance efforts. This approach requires collaboration between modelers, policy makers, and on the ground conservationists, and represents an interdisciplinary avenue for future conservation efforts.

In summary, by applying new statistical approaches for dealing with missing data to investigate the drivers of extinction more fully and considering the activities that influence threat status, we have shown that multiple traits contribute to primate extinction risk. Our findings suggest that the effects of some traits, such as home range size, have weakened over the past 20 years, indicating that the traits that influence a species’ threat status are changing as anthropogenic effects continue to transform natural landscapes. Other characteristics shown to affect extinction risk in other clades, such as behavioural flexibility, do not appear to affect primate extinction risk, suggesting that different processes likely govern extinction in different clades. Focusing on mitigating key threats, as identified here, from susceptible species’ geographic ranges will be an important and necessary step for future recovery.

## ACKNOWLEDGEMENTS

We thank Dan Greenberg, Mark Janko, and Jarrod Hadfield for statistical advice and Cecile Ane for informative conversation about Rphylopars. We also thank Susan Alberts, members of the Alberts Lab at Duke University, and Brian Lerch for their comments on previous drafts of the manuscript.

## REFERENCES

Atwood, T. B., Valentine, S. A., Hammill, E., McCauley, D. J., Madin, E. M., Beard, K. H., & Pearse, W. D. (2020). Herbivores at the highest risk of extinction among mammals, birds, and reptiles. Science Advances, 6, eabb8458.

Beever, E. A., Hall, L. E., Varner, J., Loosen, A. E., Dunham, J. B., Gahl, M. K., … & Lawler, J. J. (2017). Behavioral flexibility as a mechanism for coping with climate change. Frontiers in Ecology and the Environment, 15, 299–308.

Bennett, P. M., & Owens, I. P. (1997). Variation in extinction risk among birds: chance or evolutionary predisposition? Proceedings of the Royal Society of London. Series B: Biological Sciences, 264, 401–408.

Biber, E. (2002). Patterns of endemic extinctions among island bird species. Ecography, 25, 661–676.

Blackburn, T. M., Cassey, P., Duncan, R. P., Evans, K. L., & Gaston, K. J. (2004). Avian extinction and mammalian introductions on oceanic islands. Science, 305, 1955–1958.

Boyles, J. G., & Storm, J. J. (2007). The perils of picky eating: dietary breadth is related to extinction risk in insectivorous bats. PLoS One, 2, e672.

Butchart, S. H., Resit Akçakaya, H., Chanson, J., Baillie, J. E., Collen, B., Quader, S., … & Hilton-Taylor, C. (2007). Improvements to the red list index. PloS One, 2, e140.

Cardillo, M. (2021). Clarifying the relationship between body size and extinction risk in amphibians by complete mapping of model space. Proceedings of the Royal Society B: Biological Sciences, 288, 20203011.

Cardillo, M., & Bromham, L. (2001). Body size and risk of extinction in Australian mammals. Conservation Biology, 15, 1435–1440.

Cardillo, M., Mace, G. M., Gittleman, J. L., Jones, K. E., Bielby, J., & Purvis, A. (2008). The predictability of extinction: biological and external correlates of decline in mammals. Proceedings of the Royal Society B: Biological Sciences, 275, 1441–1448.

Cardillo, M., Mace, G. M., Jones, K. E., Bielby, J., Bininda-Emonds, O. R., Sechrest, W., … & Purvis, A. (2005). Multiple causes of high extinction risk in large mammal species. Science, 309, 1239–1241.

Cardillo, M., & Meijaard, E. (2012). Are comparative studies of extinction risk useful for conservation? Trends in Ecology & Evolution, 27, 167–171.

Chichorro, F., Correia, L., & Cardoso, P. (2022a). Biological traits interact with human threats to drive extinctions: A modelling study. Ecological Informatics, 69, 101604.

Chichorro, F., Juslén, A., & Cardoso, P. (2019). A review of the relation between species traits and extinction risk. Biological Conservation, 237, 220–229.

Chichorro, F., Urbano, F., Teixeira, D., Väre, H., Pinto, T., Brummitt, N., … & Cardoso, P. (2022b). Trait-based prediction of extinction risk across terrestrial taxa. Biological Conservation, 274, 109738.

Creighton, M. J., Greenberg, D. A., Reader, S. M., & Mooers, A. Ø. (2021). The role of behavioural flexibility in primate diversification. Animal Behaviour, 180, 269–290.

Creighton, M. J., Luo, A. Q., Reader, S. M., & Mooers, A. Ø. (2022). Predictors of taxonomic inflation and its role in primate conservation. Animal Conservation.

Davidson, A. D., Boyer, A. G., Kim, H., Pompa-Mansilla, S., Hamilton, M. J., Costa, D. P., … & Brown, J. H. (2012). Drivers and hotspots of extinction risk in marine mammals. Proceedings of the National Academy of Sciences, 109, 3395–3400.

Davidson, A. D., Hamilton, M. J., Boyer, A. G., Brown, J. H., & Ceballos, G. (2009). Multiple ecological pathways to extinction in mammals. Proceedings of the National Academy of Sciences, 106, 10702–10705.

DeCasien, A. R., Williams, S. A., & Higham, J. P. (2017). Primate brain size is predicted by diet but not sociality. Nature Ecology & Evolution, 1, 1–7.

Di Marco, M., Collen, B., Rondinini, C., & Mace, G. M. (2015). Historical drivers of extinction risk: using past evidence to direct future monitoring. Proceedings of the Royal Society B: Biological Sciences, 282, 20150928.

Ducatez, S., Sol, D., Sayol, F., & Lefebvre, L. (2020). Behavioural plasticity is associated with reduced extinction risk in birds. Nature Ecology & Evolution, 4, 788–793.

Estrada, A., Garber, P. A., Rylands, A. B., Roos, C., Fernandez-Duque, E., Di Fiore, A., … & Li, B. (2017). Impending extinction crisis of the world’s primates: Why primates matter. Science Advances, 3, e1600946.

Fox, J., Weisberg, S., Adler, D., Bates, D., Baud-Bovy, G., Ellison, S., … & Monette, G. (2012). Package ‘car’. Retrieved from: https://cran.r-project.org/web/packages/car/index.html.

Freckleton, R. P. (2002). On the misuse of residuals in ecology: regression of residuals vs. multiple regression. Journal of Animal Ecology, 71, 542–545.

Fritz, S. A., Bininda-Emonds, O. R., & Purvis, A. (2009). Geographical variation in predictors of mammalian extinction risk: big is bad, but only in the tropics. Ecology Letters, 12, 538–549.

Galán-Acedo, C., Arroyo-Rodríguez, V., Andresen, E., & Arasa-Gisbert, R. (2019). Ecological traits of the world’s primates. Scientific Data, 6, 1–5.

Gelman, A. (2008). Scaling regression inputs by dividing by two standard deviations. Statistics in Medicine, 27, 2865–2873.

González-Suárez, M., Gómez, A., & Revilla, E. (2013). Which intrinsic traits predict vulnerability to extinction depends on the actual threatening processes. Ecosphere, 4, 1–16.

Goolsby, E. W., Bruggeman, J., & Ané, C. (2017). Rphylopars: fast multivariate phylogenetic comparative methods for missing data and within-species variation. Methods in Ecology and Evolution, 8, 22–27.

Hadfield, J. D. (2010). MCMC methods for multi-response generalized linear mixed models: the MCMCglmm R package. Journal of Statistical Software. 33, 1–22.

Hadfield, J. D. (2018). MCMCglmm course notes.https://mran.microsoft.com/snapshot/2018-08-24/web/packages/MCMCglmm/vignettes/CourseNotes.pdf

Hahn, N. E., Proulx, D., Muruthi, P. M., Alberts, S., & Altmann, J. (2003). Gastrointestinal parasites in free-ranging Kenyan baboons (Papio cynocephalus and P. anubis). International Journal of Primatology, 24, 271–279.

Harcourt, A. H., & Parks, S. A. (2003). Threatened primates experience high human densities: adding an index of threat to the IUCN Red List criteria. Biological Conservation, 109, 137–149.

Hockings, K. J., Anderson, J. R., & Matsuzawa, T. (2009). Use of wild and cultivated foods by chimpanzees at Bossou, Republic of Guinea: feeding dynamics in a human-influenced environment. American Journal of Primatology, 71, 636–646.

Höglund, J. (1996). Can mating systems affect local extinction risks? Two examples of lek-breeding waders. Oikos, 77, 184–188.

IUCN. (2021). The IUCN Red List of Threatened Species Version 2021-1. IUCN Red List of Threatened Species. Retrieved from: https://www.iucnredlist.org/en.

Johnson, T. F., Isaac, N. J., Paviolo, A., & González-Suárez, M. (2021). Handling missing values in trait data. Global Ecology and Biogeography, 30, 51–62.

Leclerc, C., Courchamp, F., & Bellard, C. (2018). Insular threat associations within taxa worldwide. Scientific Reports, 8, 1–8.

Lee, T. M., & Jetz, W. (2011). Unravelling the structure of species extinction risk for predictive conservation science. Proceedings of the Royal Society B: Biological Sciences, 278, 1329–1338.

Machado, F. F., Jardim, L., Dinnage, R., Brito, D., & Cardillo, M. (2022). Diet disparity and diversity predict extinction risk in primates. Animal Conservation.

Maples, W. R., Maples, M. K., Greenhood, W. F., & Walek, M. L. (1976). Adaptations of crop-raiding baboons in Kenya. American Journal of Physical Anthropology, 45, 309–315.

Matthews, L. J., Arnold, C., Machanda, Z., & Nunn, C. L. (2011). Primate extinction risk and historical patterns of speciation and extinction in relation to body mass. Proceedings of the Royal Society B: Biological Sciences, 278, 1256–1263.

McElreath, R. (2018). Statistical rethinking: A Bayesian course with examples in R and Stan. Chapman and Hall/CRC, New York.

McLennan, M. R., Spagnoletti, N., & Hockings, K. J. (2017). The implications of primate behavioral flexibility for sustainable human–primate coexistence in anthropogenic habitats. International Journal of Primatology, 38, 105–121.

Munstermann, M. J., Heim, N. A., McCauley, D. J., Payne, J. L., Upham, N. S., Wang, S. C., & Knope, M. L. (2022). A global ecological signal of extinction risk in terrestrial vertebrates. Conservation Biology, 36, e13852.

Murray, K. A., Verde Arregoitia, L. D., Davidson, A., Di Marco, M., & Di Fonzo, M. M. (2014). Threat to the point: improving the value of comparative extinction risk analysis for conservation action. Global Change Biology, 20, 483–494.

Nakagawa, S., & De Villemereuil, P. (2019). A general method for simultaneously accounting for phylogenetic and species sampling uncertainty via Rubin’s rules in comparative analysis. Systematic Biology, 68, 632–641.

Nakagawa, S., & Freckleton, R. P. (2008). Missing inaction: the dangers of ignoring missing data. Trends in Ecology & Evolution, 23, 592–596.

Navarrete, A. F., Reader, S. M., Street, S. E., Whalen, A., & Laland, K. N. (2016). The coevolution of innovation and technical intelligence in primates. Philosophical Transactions of the Royal Society B: Biological Sciences, 371, 20150186.

Nolte, D., Boutaud, E., Kotze, D. J., Schuldt, A., & Assmann, T. (2019). Habitat specialization, distribution range size and body size drive extinction risk in carabid beetles. Biodiversity and Conservation, 28, 1267–1283.

Powell, L. E., Isler, K., & Barton, R. A. (2017). Re-evaluating the link between brain size and behavioural ecology in primates. Proceedings of the Royal Society B: Biological Sciences, 284, 20171765.

Purvis, A., Cardillo, M., Grenyer, R., & Collen, B. (2005). Correlates of extinction risk: phylogeny, biology, threat and scale. In Purvis, A., Gittleman, J.L., & Brooks, T. (Eds.), Phylogeny and Conservation (pp. 295-316). Cambridge University Press, Cambridge.

Purvis, A., Gittleman, J. L., Cowlishaw, G., & Mace, G. M. (2000). Predicting extinction risk in declining species. Proceedings of the Royal Society B: Biological Sciences, 267, 1947–1952.

Reader, S. M., Hager, Y., & Laland, K. N. (2011). The evolution of primate general and cultural intelligence. Philosophical Transactions of the Royal Society B: Biological Sciences, 366, 1017–1027.

Reader, S. M., & Laland, K. N. (2002). Social intelligence, innovation, and enhanced brain size in primates. Proceedings of the National Academy of Sciences, 99, 4436–4441.

Richards, C., Cooke, R. S., & Bates, A. E. (2021). Biological traits of seabirds predict extinction risk and vulnerability to anthropogenic threats. Global Ecology and Biogeography, 30, 973–986.

Ripley, B., Venables, B., Bates, D. M., Hornik, K., Gebhardt, A., Firth, D., & Ripley, M. B. (2013). Package ‘MASS’. CRAN Repository, Seehttp://cran.r-projectorg/web/packages/MASS/MASS.pdf.

Ripple, W. J., Wolf, C., Newsome, T. M., Hoffmann, M., Wirsing, A. J., & McCauley, D. J. (2017). Extinction risk is most acute for the world’s largest and smallest vertebrates. Proceedings of the National Academy of Sciences, 114, 10678–10683.

Rowe, N., & Myers, M. (2011). All the world’s primates. Primate Conservation, Inc. Retrieved from: https://www.alltheworldsprimates.org.

Rueda-Cediel, P., Anderson, K. E., Regan, T. J., & Regan, H. M. (2018). Effects of uncertainty and variability on population declines and IUCN Red List classifications. Conservation Biology, 32, 916–925.

Ruland, F., & Jeschke, J. M. (2017). Threat-dependent traits of endangered frogs. Biological Conservation, 206, 310–313.

Rylands, A. B., & Mittermeier, R. A. (2014). Primate taxonomy: species and conservation. Evolutionary Anthropology, 23, 8–10.

Salafsky, N., Salzer, D., Stattersfield, A. J., Hilton-Taylor, C. R. A. I. G., Neugarten, R., Butchart, S. H., … & Wilkie, D. (2008). A standard lexicon for biodiversity conservation: unified classifications of threats and actions. Conservation Biology, 22, 897–911.

Santos, T., Diniz-Filho, J. A., e Luis, T. R., Bini, M., & Santos, M. T. (2018). Package ‘PVR’. Retrieved from: https://cran.r-project.org/web/packages/PVR/index.html.

Upham, N. S., Esselstyn, J. A., & Jetz, W. (2019). Inferring the mammal tree: Species-level sets of phylogenies for questions in ecology, evolution, and conservation. PLoS Biology, 17, 1–44.

Woodroffe, R., & Ginsberg, J. R. (1998). Edge effects and the extinction of populations inside protected areas. Science, 280, 2126–2128.

