## Supplementary Materials for "Explaining the primate extinction crisis: predictors of extinction risk and active threats"

### SUPPLEMENTARY METHODS:

#### DATA:

Data on body mass (kg) were taken from the compilation by Galán-Acedo et al. (2019) and converted to grams. In cases where multiple sources of body mass data were available from Galán-Acedo et al. (2019), we took data from the source where female body mass was available. If either multiple sources or no sources had data available on female body mass, we preferred sources with data on wild populations and the largest sample sizes. Data on home range size (ha) were also taken from Galán-Acedo et al. (2019). In cases where multiple sources of home range size were available from Galán-Acedo et al. (2019) we took data from the source with the greatest sample size and those with the longest study duration. We also preferred sources that made this information and their methodology available. We did not average trait values from multiple sources in Galán-Acedo et al. (2019) because there was uncertainty as to whether there was overlap in sampled populations in different compilations/sources.

Generation length estimates were obtained from the International Union for Conservation of Nature (IUCN, 2021). Generation lengths that were given as a range by the IUCN were taken as the midpoint between the lower and upper range values (e.g., generation lengths of 6 to 7 years taken as 6.5 years). One species with data from the IUCN, *Pithecia vanzolinii*, had an abnormally large generation length which we deemed to be incorrect and thus did not use in our dataset.

We surveyed all references in DeCasien et al. (2017) to supplement group size data that were available for named species not included in their species list and created averages for these species following DeCasien et al. (2017). For species in our data set that were still missing group size data, we then surveyed Rowe & Myers (2011) for data on average group size. In cases where Rowe & Myers (2011) had multiple reports for average group size, we took the geometric mean of averages provided from different populations. We did not include measures of average group size reported in Rowe & Myers

(2011) that only included part of a larger group (e.g., size of family units in species that spend most of their time in larger groups).

Data on social system (solitary, pair-living, harem polygyny and polygynandry) were also taken from Decasien et al. (2017) and supplemented with information from Rowe & Myers (2011). In cases where neither Decasien et al. (2017) or Rowe & Myers (2011) provided references for social system data, we filled in data gaps by conducting our own literature review. In cases where relatively new or understudied species had no information published on the species' social system, but the entire genus was characterized by one dominant social system, we accepted the dominant social system as the species system. While there is variability in social organization within species, we chose the predominant social system for each species according to available literature.

Insularity was inferred from range data available from the Rowe & Myers (2011) and the IUCN (2021). Species insularity was coded as 'true' for insular species only occurring on islands (i.e., island endemics) and 'false' for all other species. Species found exclusively on Madagascar, Borneo, or Sumatra were not scored as insular since these islands are large enough to support large geographic ranges comparable to many mainland species. Lifestyle (arboreal, terrestrial, or both) was recorded from Rowe & Myers (2011) and, when unavailable from Rowe & Myers (2011), was filled in using accounts from the literature by searching species name and "arboreal" and species name and "terrestrial" (see full list of references in the Supplementary Data). Nocturnal (true or false) was recorded from Estrada et al. (2017) and information provided by the IUCN (2021).

ANALYSIS:

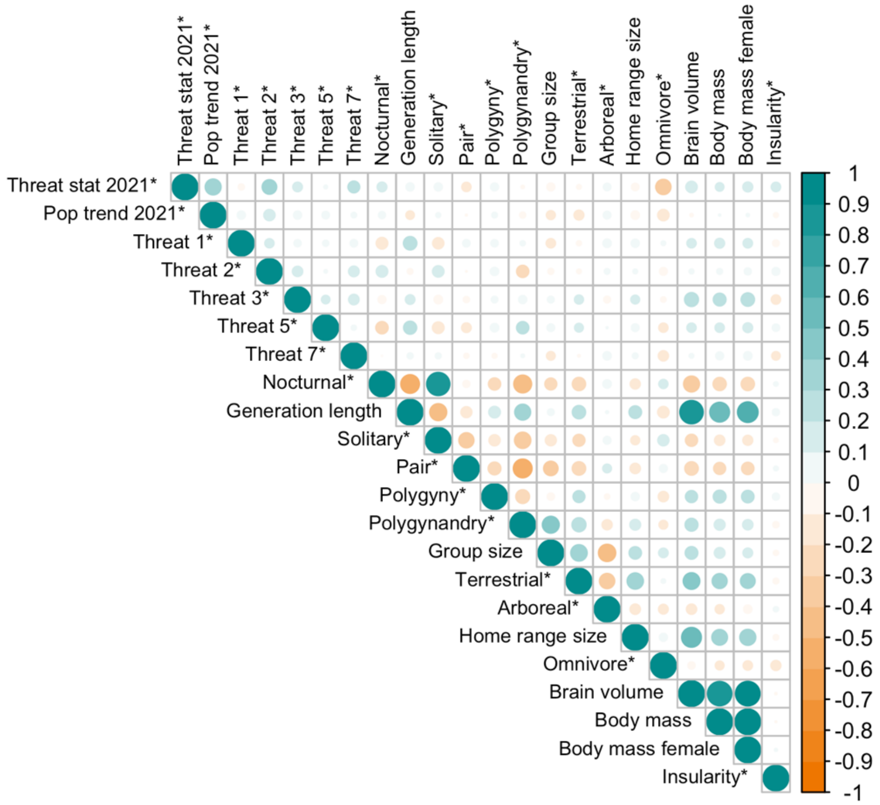


Figure S1: Correlation matrix of primate threat status, population trend, threats, and possible predictors of primate extinction risk for 446 primate species. *Marks variables that are scored as binary (0/1).

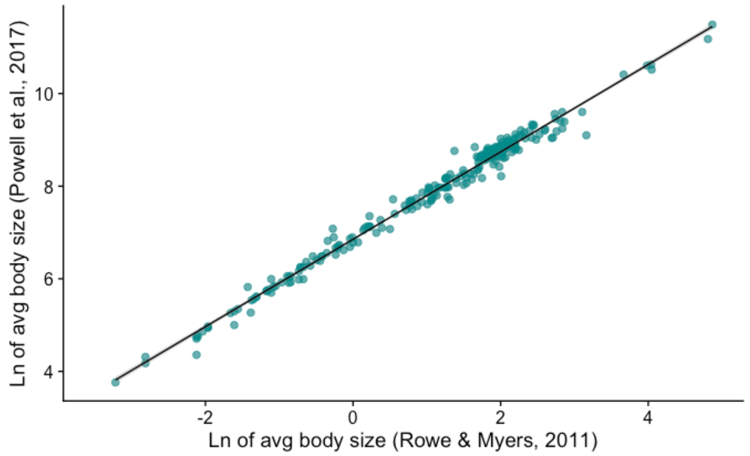


Figure S2: Body mass data from Powell et al. (2017) versus Rowe & Myers (2011) (r=0.98).

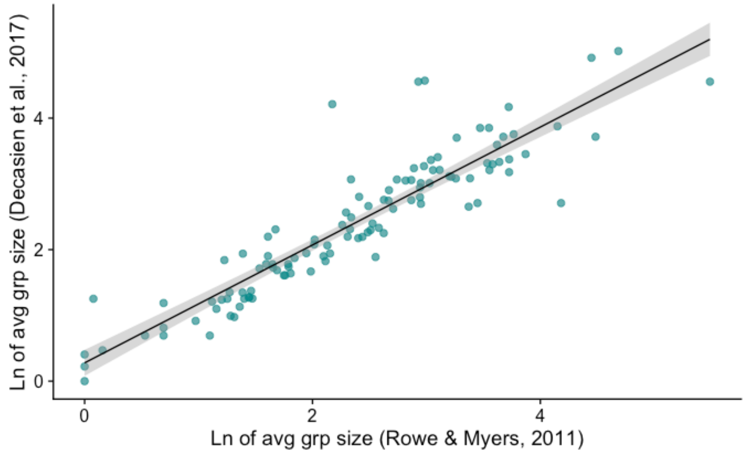


Figure S3: Group size data from DeCasien et al. (2017) versus Rowe & Myers (2011) (r=0.83).

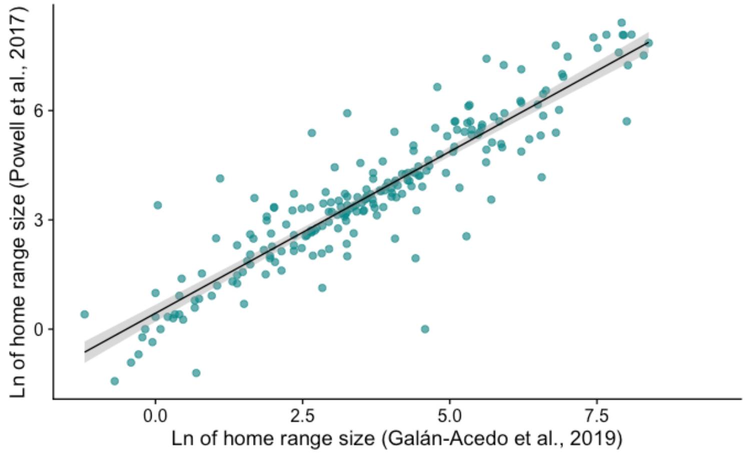


Figure S4: Home range size data from Powell et al. (2017) versus Galán-Acedo et al. (2019) (r=0.69).

Table S1: Available data (%) and average predictive accuracy ($p^2$) or average area under the ROC curve (AUC) scores across 100 imputations for predictor variables (n=446), with the number of eigenvectors included in imputation models.

| **Variable (Units)** | **# Species with available data (%)** | **# of eigenvectors** | **Average predictive accuracy ($p^2$) or area under the ROC curve (AUC)** |
|---|---|---|---|
| **Body mass (g)** | 419 (94%) | 20 | $p^2$=0.912 |
| **Female body mass (g)** | 317 (72%) | 60 | $p^2$=0.950 |
| **Omnivory (true or false)** | 372 (83%) | 60 | AUC= 0.945 |
| **Generation length (yrs)** | 343 (77%) | 10 | $p^2$=0.926 |
| **Home range size (ha)** | 333 (75%) | 30 | $p^2$=0.993 |
| **Group size** | 271 (61%) | 30 | $p^2$=0.903 |
| **Brain volume ($cm^3$)** | 241 (54%) | 30 | $p^2$=0.986 |

### SUPPLEMENTARY RESULTS:

*Predictors of threat status and population trends*

Table S2: Predictors of threat status (ordinal; LC=0, NT=1, VU=2, EN=3, and CR=4) for 430 species with known threat statuses. Posterior means and standard error pooled from 100 MCMCglmm models (one per imputed dataset and sampled phylogeny) with ordinal error structures. Percentage of positive and negative coefficients come from 1,001 iterations retained across each of the 100 models (total retained iterations = 100,100). Each model included 10 traits as fixed effects and continuous variables ‡ were ln-transformed, centered with respect to the mean, and scaled by 2 standard deviations.

| term | %coef + | %coef - | pooled pm | pooled std.error |
|---|---|---|---|---|
| **(Intercept)** | 91 | 9 | 1.572 | 1.180 |
| **Body mass (g) ‡** | *93* | *7* | *0.980* | *0.648* |
| **Generation length ‡** | 71 | 29 | 0.237 | 0.430 |
| **Home range size (ha) ‡** | 33 | 67 | -0.149 | 0.345 |
| **Group size ‡** | *6* | *94* | *-0.561* | *0.359* |
| **Brain volume ($cm^3$) ‡** | 56 | 44 | 0.074 | 0.476 |
| **Omnivore** | *5* | *95* | *-0.474* | *0.293* |
| **Social system: Pair living** | 42 | 58 | -0.129 | 0.607 |
| **Social system: Polygynandry** | 43 | 57 | -0.125 | 0.674 |
| **Social system: Polygyny** | 79 | 21 | 0.565 | 0.728 |
| **Lifestyle: Arboreal** | 43 | 57 | -0.097 | 0.533 |
| **Lifestyle: Both** | 55 | 45 | 0.066 | 0.512 |
| **Insularity** | *100* | *0* | *1.214* | *0.428* |
| **Nocturnal** | 21 | 79 | -0.535 | 0.664 |

Table S3: Predictors of threat status (ordinal; LC or NT=0, VU=1, EN=2, and CR=3) for 208 species with known threat statuses in Harcourt & Parks (2003). Posterior means and standard error pooled from 100 MCMCglmm models (one per imputed dataset and sampled phylogeny) with ordinal error structures. Percentage of positive and negative coefficients come from 1,001 iterations retained across each of the 100 models (total retained iterations = 100,100). Each model included 10 traits as fixed effects and continuous variables ‡ were ln-transformed, centered with respect to the mean, and scaled by 2 standard deviations.

| **term** | **%coef +** | **%coef -** | **pooled pm** | **pooled std.error** |
|---|---|---|---|---|
| **(Intercept)** | 38 | 62 | -0.415 | 1.390 |
| **Body mass (g) ‡** | 84 | 16 | 0.934 | 0.955 |
| **Generation length ‡** | 50 | 50 | 0.006 | 0.669 |
| **Home range size (ha) ‡** | *100* | *0* | *1.498* | *0.562* |
| **Group size ‡** | 23 | 77 | -0.443 | 0.610 |
| **Brain volume ($cm^3$) ‡** | 62 | 38 | 0.327 | 1.094 |
| **Omnivore** | 15 | 85 | -0.499 | 0.480 |
| **Social system: Pair living** | 62 | 38 | 0.224 | 0.773 |
| **Social system: Polygynandry** | 38 | 62 | -0.254 | 0.796 |
| **Social system: Polygyny** | 66 | 34 | 0.340 | 0.852 |
| **Lifestyle: Arboreal** | 65 | 35 | 0.287 | 0.740 |
| **Lifestyle: Both** | 51 | 49 | 0.015 | 0.713 |
| **Insularity** | *100* | *0* | *2.255* | *0.661* |
| **Nocturnal** | 29 | 71 | -0.464 | 0.854 |

Table S4: Predictors of threat status (binary) for 430 species with known threat statuses. Posterior means and standard error pooled from 100 MCMCglmm models (one per imputed dataset and sampled phylogeny) with threshold error structures. Percentage of positive and negative coefficients come from 1,001 iterations retained across each of the 100 models (total retained iterations = 100,100). Each model included 10 traits as fixed effects and continuous variables ‡ were ln-transformed, centered with respect to the mean, and scaled by 2 standard deviations.

| term | %coef + | %coef - | pooled pm | pooled std.error |
|---|---|---|---|---|
| **(Intercept)** | 56 | 44 | 0.193 | 1.407 |
| **Body mass (g) ‡** | 77 | 23 | 0.629 | 0.875 |
| **Generation length ‡** | 66 | 34 | 0.235 | 0.601 |
| **Home range size (ha) ‡** | 37 | 63 | -0.151 | 0.481 |
| **Group size ‡** | 9 | 91 | -0.665 | 0.538 |
| **Brain volume ($cm^3$) ‡** | 55 | 45 | 0.066 | 0.649 |
| **Omnivore** | 18 | 82 | -0.314 | 0.363 |
| **Social system: Pair living** | 75 | 25 | 0.599 | 0.899 |
| **Social system: Polygynandry** | 58 | 42 | 0.169 | 0.874 |
| **Social system: Polygyny** | 85 | 15 | 0.949 | 0.899 |
| **Lifestyle: Arboreal** | 67 | 33 | 0.244 | 0.590 |
| **Lifestyle: Both** | 64 | 36 | 0.196 | 0.562 |
| **Insularity** | 86 | 14 | 0.746 | 0.720 |
| **Nocturnal** | 57 | 43 | 0.151 | 0.915 |

Table S5: Results from tests of body mass, generation length, and brain (separate models) versus threat status (binary) without other fixed effects for 430 species with known threat statuses. Posterior means and standard error pooled from 100 MCMCglmm models (one per imputed dataset and sampled phylogeny) with threshold error structures. Percentage of positive and negative coefficients come from 1,001 iterations retained across each of the 100 models (total retained iterations = 100,100). Traits were included as fixed effects in separate models and continuous variables ‡ were ln-transformed, centered with respect to the mean, and scaled by 2 standard deviations.

| **term** | **%coef +** | **%coef -** | **pooled pm** | **pooled std.error** |
|---|---|---|---|---|
| **(Intercept)** | 81 | 19 | 0.871 | 0.994 |
| **Body mass (g) ‡** | 78 | 22 | 0.462 | 0.618 |
| **(Intercept)** | 69 | 31 | 0.560 | 1.173 |
| **Generation length ‡** | 75 | 25 | 0.321 | 0.484 |
| **(Intercept)** | 68 | 32 | 0.560 | 1.183 |
| **Brain volume ($cm^3$) ‡** | 60 | 40 | 0.121 | 0.525 |

Table S6: Predictors of threat status (binary) for 208 species with known threat statuses in Harcourt & Parks (2003). Posterior means and standard error pooled from 100 MCMCglmm models (one per imputed dataset and sampled phylogeny) with threshold error structures. Percentage of positive and negative coefficients come from 1,001 iterations retained across each of the 100 models (total retained iterations = 100,100). Each model included 10 traits as fixed effects and continuous variables ‡ were ln-transformed, centered with respect to the mean, and scaled by 2 standard deviations.

| **term** | **%coef +** | **%coef -** | **pooled pm** | **pooled std.error** |
|---|---|---|---|---|
| **(Intercept)** | 40 | 60 | -0.295 | 1.193 |
| **Body mass (g) ‡** | 94 | 6 | 1.276 | 0.847 |
| **Generation length ‡** | 41 | 59 | -0.116 | 0.543 |
| **Home range size (ha) ‡** | *99* | *1* | *1.078* | *0.466* |
| **Group size ‡** | 20 | 80 | -0.433 | 0.527 |
| **Brain volume ($cm^3$) ‡** | 60 | 40 | 0.243 | 0.955 |
| **Omnivore** | 18 | 82 | -0.335 | 0.384 |
| **Social system: Pair living** | 58 | 42 | 0.138 | 0.695 |
| **Social system: Polygynandry** | 25 | 75 | -0.486 | 0.746 |
| **Social system: Polygyny** | 57 | 43 | 0.129 | 0.794 |
| **Lifestyle: Arboreal** | 73 | 27 | 0.380 | 0.637 |
| **Lifestyle: Both** | 61 | 39 | 0.170 | 0.604 |
| **Insularity** | *100* | *0* | *2.556* | *0.810* |
| **Nocturnal** | 28 | 72 | -0.415 | 0.722 |

Table S7: Predictors of population trend (binary) for 401 species with known population trends. Posterior means and standard error pooled from 100 MCMCglmm models (one per imputed dataset and sampled phylogeny) with threshold error structures. Percentage of positive and negative coefficients come from 1,001 iterations retained across each of the 100 models (total retained iterations = 100,100). Each model included 10 traits as fixed effects and continuous variables ‡ were ln-transformed, centered with respect to the mean, and scaled by 2 standard deviations.

| term | %coef + | %coef - | pooled pm | pooled std.error |
|---|---|---|---|---|
| **(Intercept)** | 88 | 12 | 1.522 | 1.319 |
| **Body mass (g) ‡** | 53 | 47 | 0.075 | 0.912 |
| **Generation length ‡** | 44 | 56 | -0.110 | 0.751 |
| **Home range size (ha) ‡** | 15 | 85 | -0.530 | 0.558 |
| **Group size ‡** | 10 | 90 | -0.758 | 0.643 |
| **Brain volume ($cm^3$) ‡** | 52 | 48 | 0.044 | 0.729 |
| **Omnivore** | 5 | 95 | -0.698 | 0.480 |
| **Social system: Pair living** | 39 | 61 | -0.228 | 0.873 |
| **Social system: Polygynandry** | 61 | 39 | 0.228 | 0.847 |
| **Social system: Polygyny** | 69 | 31 | 0.428 | 0.882 |
| **Lifestyle: Arboreal** | 66 | 34 | 0.251 | 0.626 |
| **Lifestyle: Both** | 42 | 58 | -0.115 | 0.581 |
| **Insularity** | 10 | 90 | -0.792 | 0.643 |
| **Nocturnal** | 73 | 27 | 0.608 | 1.024 |

Table S8: Predictors of threat status (ordinal; LC=0, NT=1, VU=2, EN=3, and CR=4) for 430 species with known threat statuses using imputed traits values from Rphylopars (Goolsby et al., 2017). Posterior means and standard error pooled from 100 MCMCglmm models (one per imputed dataset and sampled phylogeny) with ordinal error structures. Percentage of positive and negative coefficients come from 1,001 iterations retained across each of the 100 models (total retained iterations = 100,100). Each model included 10 traits as fixed effects and continuous variables ‡ were ln-transformed, centered with respect to the mean, and scaled by 2 standard deviations.

| term | %coef + | %coef - | pooled pm | pooled std.error |
|---|---|---|---|---|
| **(Intercept)** | 91 | 9 | 1.639 | 1.185 |
| **Body mass (g) ‡** | 74 | 26 | 0.517 | 0.828 |
| **Generation length ‡** | 86 | 14 | 0.485 | 0.461 |
| **Home range size (ha) ‡** | 29 | 71 | -0.204 | 0.359 |
| **Group size ‡** | *2* | *98* | *-0.831* | *0.413* |
| **Brain volume ($cm^3$) ‡** | 79 | 21 | 0.857 | 1.058 |
| **Omnivore** | *9* | *91* | *-0.391* | *0.296* |
| **Social system: Pair living** | 43 | 57 | -0.106 | 0.607 |
| **Social system: Polygynandry** | 44 | 56 | -0.108 | 0.684 |
| **Social system: Polygyny** | 74 | 26 | 0.463 | 0.731 |
| **Lifestyle: Arboreal** | 37 | 63 | -0.178 | 0.534 |
| **Lifestyle: Both** | 49 | 51 | -0.008 | 0.511 |
| **Insularity** | *100* | *0* | *1.225* | *0.428* |
| **Nocturnal** | 22 | 78 | -0.512 | 0.668 |

Table S9: Predictors of threat status (binary) for 430 species with known threat statuses using imputed traits values from Rphylopars (Goolsby et al., 2017). Posterior means and standard error pooled from 100 MCMCglmm models (one per imputed dataset and sampled phylogeny) with threshold error structures. Percentage of positive and negative coefficients come from 1,001 iterations retained across each of the 100 models (total retained iterations = 100,100). Each model included 10 traits as fixed effects and continuous variables ‡ were ln-transformed, centered with respect to the mean, and scaled by 2 standard deviations.

| term | %coef + | %coef - | pooled pm | pooled std.error |
|---|---|---|---|---|
| **(Intercept)** | 58 | 42 | 0.286 | 1.383 |
| **Body mass (g) ‡** | 68 | 32 | 0.434 | 0.951 |
| **Generation length ‡** | 90 | 10 | 0.842 | 0.673 |
| **Home range size (ha) ‡** | 38 | 62 | -0.137 | 0.454 |
| **Group size ‡** | 5 | 95 | -0.848 | 0.549 |
| **Brain volume ($cm^3$) ‡** | 61 | 39 | 0.324 | 1.158 |
| **Omnivore** | 24 | 76 | -0.242 | 0.344 |
| **Social system: Pair living** | 79 | 21 | 0.683 | 0.876 |
| **Social system: Polygynandry** | 58 | 42 | 0.162 | 0.874 |
| **Social system: Polygyny** | 82 | 18 | 0.808 | 0.901 |
| **Lifestyle: Arboreal** | 62 | 38 | 0.166 | 0.574 |
| **Lifestyle: Both** | 59 | 41 | 0.124 | 0.543 |
| **Insularity** | 87 | 13 | 0.733 | 0.701 |
| **Nocturnal** | 56 | 44 | 0.136 | 0.893 |

Table S10: Predictors of population trend (binary) for 401 species with known population trends using imputed traits values from Rphylopars (Goolsby et al., 2017). Posterior means and standard error pooled from 100 MCMCglmm models (one per imputed dataset and sampled phylogeny) with threshold error structures. Percentage of positive and negative coefficients come from 1,001 iterations retained across each of the 100 models (total retained iterations = 100,100). Each model included 10 traits as fixed effects and continuous variables ‡ were ln-transformed, centered with respect to the mean, and scaled by 2 standard deviations.

| **term** | **%coef +** | **%coef -** | **pooled pm** | **pooled std.error** |
|---|---|---|---|---|
| **(Intercept)** | 89 | 11 | 1.579 | 1.318 |
| **Body mass (g) ‡** | 51 | 49 | 0.026 | 0.955 |
| **Generation length ‡** | 50 | 50 | -0.001 | 0.803 |
| **Home range size (ha) ‡** | 12 | 88 | -0.602 | 0.551 |
| **Group size ‡** | 6 | 94 | -0.836 | 0.590 |
| **Brain volume ($cm^3$) ‡** | 51 | 49 | 0.035 | 1.169 |
| **Omnivore** | 4 | 96 | -0.757 | 0.478 |
| **Social system: Pair living** | 38 | 62 | -0.260 | 0.879 |
| **Social system: Polygynandry** | 64 | 36 | 0.309 | 0.853 |
| **Social system: Polygyny** | 67 | 33 | 0.375 | 0.886 |
| **Lifestyle: Arboreal** | 61 | 39 | 0.171 | 0.624 |
| **Lifestyle: Both** | 41 | 59 | -0.134 | 0.578 |
| **Insularity** | 9 | 91 | -0.825 | 0.632 |
| **Nocturnal** | 72 | 28 | 0.573 | 1.028 |

*Predictors of specific threat types*

Table S11: Predictors of threat type 1: residential and commercial development (binary) for 404 species with known threats. Posterior means and standard error pooled from 100 MCMCglmm models (one per imputed dataset and sampled phylogeny) with threshold error structures. Percentage of positive and negative coefficients come from 1,001 iterations retained across each of the 100 models (total retained iterations = 100,100). Each model included 10 traits as fixed effects and continuous variables ‡ were ln-transformed, centered with respect to the mean, and scaled by 2 standard deviations.

| term | %coef + | %coef - | pooled pm | pooled std.error |
|---|---|---|---|---|
| **(Intercept)** | 7 | 93 | -1.494 | 1.003 |
| **Body mass (g) ‡** | *96* | *4* | *1.004* | *0.591* |
| **Generation length ‡** | 63 | 37 | 0.143 | 0.422 |
| **Home range size (ha) ‡** | 30 | 70 | -0.162 | 0.320 |
| **Group size ‡** | 13 | 87 | -0.400 | 0.368 |
| **Brain volume ($cm^3$) ‡** | 56 | 44 | 0.058 | 0.433 |
| **Omnivore** | 76 | 24 | 0.185 | 0.258 |
| **Social system: Pair living** | 23 | 77 | -0.405 | 0.568 |
| **Social system: Polygynandry** | 23 | 77 | -0.473 | 0.645 |
| **Social system: Polygyny** | 15 | 85 | -0.675 | 0.651 |
| **Lifestyle: Arboreal** | *98* | *2* | *1.045* | *0.545* |
| **Lifestyle: Both** | 87 | 13 | 0.591 | 0.536 |
| **Insularity** | *2* | *98* | *-0.752* | *0.381* |
| **Nocturnal** | 87 | 13 | 0.683 | 0.625 |

Table S12: Predictors of threat type 2: agriculture and aquaculture (binary) for 404 species with known threats. Posterior means and standard error pooled from 100 MCMCglmm models (one per imputed dataset and sampled phylogeny) with threshold error structures. Percentage of positive and negative coefficients come from 1,001 iterations retained across each of the 100 models (total retained iterations = 100,100). Each model included 10 traits as fixed effects and continuous variables ‡ were ln-transformed, centered with respect to the mean, and scaled by 2 standard deviations.

| term | %coef + | %coef - | pooled pm | pooled std.error |
|---|---|---|---|---|
| **(Intercept)** | 84 | 16 | 1.524 | 1.544 |
| **Body mass (g) ‡** | 55 | 45 | 0.110 | 0.997 |
| **Generation length ‡** | 44 | 56 | -0.103 | 0.821 |
| **Home range size (ha) ‡** | 72 | 28 | 0.332 | 0.627 |
| **Group size ‡** | 44 | 56 | -0.115 | 0.691 |
| **Brain volume ($cm^3$) ‡** | 54 | 46 | 0.064 | 0.776 |
| **Omnivore** | 60 | 40 | 0.114 | 0.519 |
| **Social system: Pair living** | 35 | 65 | -0.356 | 0.966 |
| **Social system: Polygynandry** | 15 | 85 | -0.955 | 0.920 |
| **Social system: Polygyny** | 45 | 55 | -0.113 | 1.002 |
| **Lifestyle: Arboreal** | 84 | 16 | 0.686 | 0.713 |
| **Lifestyle: Both** | 66 | 34 | 0.268 | 0.671 |
| **Insularity** | 83 | 17 | 0.914 | 1.055 |
| **Nocturnal** | 48 | 52 | -0.060 | 0.936 |

Table S13: Predictors of threat type 3: energy production and mining (binary) for 404 species with known threats. Posterior means and standard error pooled from 100 MCMCglmm models (one per imputed dataset and sampled phylogeny) with threshold error structures. Percentage of positive and negative coefficients come from 1,001 iterations retained across each of the 100 models (total retained iterations = 100,100). Each model included 10 traits as fixed effects and continuous variables ‡ were ln-transformed, centered with respect to the mean, and scaled by 2 standard deviations.

| **term** | **%coef +** | **%coef -** | **pooled pm** | **pooled std.error** |
|---|---|---|---|---|
| **(Intercept)** | 10 | 90 | -1.081 | 0.821 |
| **Body mass (g) ‡** | *99* | *1* | *1.232* | *0.514* |
| **Generation length ‡** | 14 | 86 | -0.366 | 0.338 |
| **Home range size (ha) ‡** | 27 | 73 | -0.176 | 0.286 |
| **Group size ‡** | 34 | 66 | -0.143 | 0.343 |
| **Brain volume ($cm^3$) ‡** | 56 | 44 | 0.064 | 0.405 |
| **Omnivore** | 93 | 7 | 0.371 | 0.255 |
| **Social system: Pair living** | 90 | 10 | 0.545 | 0.440 |
| **Social system: Polygynandry** | 80 | 20 | 0.434 | 0.523 |
| **Social system: Polygyny** | 71 | 29 | 0.291 | 0.548 |
| **Lifestyle: Arboreal** | 48 | 52 | -0.026 | 0.468 |
| **Lifestyle: Both** | 58 | 42 | 0.091 | 0.453 |
| **Insularity** | *0* | *100* | *-1.807* | *0.793* |
| **Nocturnal** | 59 | 41 | 0.095 | 0.453 |

Table S14: Predictors of threat type 5: biological resource use (binary) for 404 species with known threats. Posterior means and standard error pooled from 100 MCMCglmm models (one per imputed dataset and sampled phylogeny) with threshold error structures. Percentage of positive and negative coefficients come from 1,001 iterations retained across each of the 100 models (total retained iterations = 100,100). Each model included 10 traits as fixed effects and continuous variables ‡ were ln-transformed, centered with respect to the mean, and scaled by 2 standard deviations.

| term | %coef + | %coef - | pooled pm | pooled std.error |
|---|---|---|---|---|
| **(Intercept)** | 85 | 15 | 1.042 | 1.057 |
| **Body mass (g) ‡** | 84 | 16 | 0.605 | 0.649 |
| **Generation length ‡** | 91 | 9 | 0.582 | 0.461 |
| **Home range size (ha) ‡** | 85 | 15 | 0.458 | 0.458 |
| **Group size ‡** | 73 | 27 | 0.316 | 0.519 |
| **Brain volume ($cm^3$) ‡** | 52 | 48 | 0.033 | 0.497 |
| **Omnivore** | 61 | 39 | 0.088 | 0.345 |
| **Social system: Pair living** | 25 | 75 | -0.351 | 0.547 |
| **Social system: Polygynandry** | 74 | 26 | 0.444 | 0.708 |
| **Social system: Polygyny** | 70 | 30 | 0.401 | 0.773 |
| **Lifestyle: Arboreal** | 87 | 13 | 0.825 | 0.714 |
| **Lifestyle: Both** | 89 | 11 | 0.900 | 0.725 |
| **Insularity** | 93 | 7 | 1.158 | 0.926 |
| **Nocturnal** | 45 | 55 | -0.078 | 0.595 |

Table S15: Predictors of threat type 7: natural system modifications (binary) for 404 species with known threats. Posterior means and standard error pooled from 100 MCMCglmm models (one per imputed dataset and sampled phylogeny) with threshold error structures. Percentage of positive and negative coefficients come from 1,001 iterations retained across each of the 100 models (total retained iterations = 100,100). Each model included 10 traits as fixed effects and continuous variables ‡ were ln-transformed, centered with respect to the mean, and scaled by 2 standard deviations.

| term | %coef + | %coef - | pooled pm | pooled std.error |
|---|---|---|---|---|
| **(Intercept)** | 24 | 76 | -0.723 | 1.031 |
| **Body mass (g) ‡** | 91 | 9 | 0.699 | 0.535 |
| **Generation length ‡** | 25 | 75 | -0.218 | 0.335 |
| **Home range size (ha) ‡** | 47 | 53 | -0.023 | 0.317 |
| **Group size ‡** | 44 | 56 | -0.055 | 0.339 |
| **Brain volume ($cm^3$) ‡** | 51 | 49 | 0.016 | 0.417 |
| **Omnivore** | 31 | 69 | -0.141 | 0.280 |
| **Social system: Pair living** | 28 | 72 | -0.255 | 0.460 |
| **Social system: Polygynandry** | 18 | 82 | -0.488 | 0.538 |
| **Social system: Polygyny** | 7 | 93 | -0.820 | 0.571 |
| **Lifestyle: Arboreal** | 92 | 8 | 0.927 | 0.709 |
| **Lifestyle: Both** | 95 | 5 | 1.047 | 0.698 |
| **Insularity** | *0* | *100* | *-1.312* | *0.533* |
| **Nocturnal** | 14 | 86 | -0.523 | 0.513 |

Table S16: Predictors of threat type 1: residential and commercial development (binary) for 404 species with known threats using imputed traits values from Rphylopars (Goolsby et al., 2017). Posterior means and standard error pooled from 100 MCMCglmm models (one per imputed dataset and sampled phylogeny) with threshold error structures. Percentage of positive and negative coefficients come from 1,001 iterations retained across each of the 100 models (total retained iterations = 100,100). Each model included 10 traits as fixed effects and continuous variables ‡ were ln-transformed, centered with respect to the mean, and scaled by 2 standard deviations.

| **term** | **%coef +** | **%coef -** | **pooled pm** | **pooled std.error** |
|---|---|---|---|---|
| **(Intercept)** | 8 | 92 | -1.439 | 1.024 |
| **Body mass (g) ‡** | 83 | 17 | 0.741 | 0.783 |
| **Generation length ‡** | 67 | 33 | 0.201 | 0.466 |
| **Home range size (ha) ‡** | 25 | 75 | -0.220 | 0.328 |
| **Group size ‡** | *3* | *97* | *-0.720* | *0.387* |
| **Brain volume ($cm^3$) ‡** | 73 | 27 | 0.619 | 1.023 |
| **Omnivore** | 74 | 26 | 0.167 | 0.261 |
| **Social system: Pair living** | 26 | 74 | -0.369 | 0.579 |
| **Social system: Polygynandry** | 29 | 71 | -0.369 | 0.666 |
| **Social system: Polygyny** | 14 | 86 | -0.720 | 0.666 |
| **Lifestyle: Arboreal** | *97* | *3* | *0.956* | *0.550* |
| **Lifestyle: Both** | 84 | 16 | 0.519 | 0.543 |
| **Insularity** | *2* | *98* | *-0.748* | *0.381* |
| **Nocturnal** | 86 | 14 | 0.672 | 0.632 |

Table S17: Predictors of threat type 2: agriculture and aquaculture (binary) for 404 species with known threats using imputed traits values from Rphylopars (Goolsby et al., 2017). Posterior means and standard error pooled from 100 MCMCglmm models (one per imputed dataset and sampled phylogeny) with threshold error structures. Percentage of positive and negative coefficients come from 1,001 iterations retained across each of the 100 models (total retained iterations = 100,100). Each model included 10 traits as fixed effects and continuous variables ‡ were ln-transformed, centered with respect to the mean, and scaled by 2 standard deviations.

| term | %coef + | %coef - | pooled pm | pooled std.error |
|---|---|---|---|---|
| **(Intercept)** | 85 | 15 | 1.608 | 1.514 |
| **Body mass (g) ‡** | 57 | 43 | 0.164 | 1.033 |
| **Generation length ‡** | 52 | 48 | 0.043 | 0.803 |
| **Home range size (ha) ‡** | 68 | 32 | 0.244 | 0.607 |
| **Group size ‡** | 51 | 49 | -0.001 | 0.667 |
| **Brain volume ($cm^3$) ‡** | 50 | 50 | 0.000 | 1.192 |
| **Omnivore** | 53 | 47 | 0.032 | 0.505 |
| **Social system: Pair living** | 35 | 65 | -0.349 | 0.943 |
| **Social system: Polygynandry** | 14 | 86 | -1.002 | 0.916 |
| **Social system: Polygyny** | 43 | 57 | -0.167 | 0.989 |
| **Lifestyle: Arboreal** | 84 | 16 | 0.672 | 0.697 |
| **Lifestyle: Both** | 66 | 34 | 0.253 | 0.662 |
| **Insularity** | 83 | 17 | 0.908 | 1.051 |
| **Nocturnal** | 48 | 52 | -0.056 | 0.908 |

Table S18: Predictors of threat type 3: energy production and mining (binary) for 404 species with known threats using imputed traits values from Rphylopars (Goolsby et al., 2017). Posterior means and standard error pooled from 100 MCMCglmm models (one per imputed dataset and sampled phylogeny) with threshold error structures. Percentage of positive and negative coefficients come from 1,001 iterations retained across each of the 100 models (total retained iterations = 100,100). Each model included 10 traits as fixed effects and continuous variables ‡ were ln-transformed, centered with respect to the mean, and scaled by 2 standard deviations.

| term | %coef + | %coef - | pooled pm | pooled std.error |
|---|---|---|---|---|
| **(Intercept)** | 9 | 91 | -1.083 | 0.817 |
| **Body mass (g) ‡** | *92* | *8* | *1.031* | *0.730* |
| **Generation length ‡** | 7 | 93 | -0.523 | 0.365 |
| **Home range size (ha) ‡** | 23 | 77 | -0.217 | 0.296 |
| **Group size ‡** | 27 | 73 | -0.206 | 0.338 |
| **Brain volume ($cm^3$) ‡** | 72 | 28 | 0.537 | 0.938 |
| **Omnivore** | 90 | 10 | 0.324 | 0.253 |
| **Social system: Pair living** | 91 | 9 | 0.568 | 0.440 |
| **Social system: Polygynandry** | 82 | 18 | 0.467 | 0.527 |
| **Social system: Polygyny** | 71 | 29 | 0.304 | 0.548 |
| **Lifestyle: Arboreal** | 46 | 54 | -0.041 | 0.467 |
| **Lifestyle: Both** | 58 | 42 | 0.088 | 0.449 |
| **Insularity** | *0* | *100* | *-1.803* | *0.793* |
| **Nocturnal** | 63 | 37 | 0.139 | 0.452 |

Table S19: Predictors of threat type 5: biological resource use (binary) for 404 species with known threats using imputed traits values from Rphylopars (Goolsby et al., 2017). Posterior means and standard error pooled from 100 MCMCglmm models (one per imputed dataset and sampled phylogeny) with threshold error structures. Percentage of positive and negative coefficients come from 1,001 iterations retained across each of the 100 models (total retained iterations = 100,100). Each model included 10 traits as fixed effects and continuous variables ‡ were ln-transformed, centered with respect to the mean, and scaled by 2 standard deviations.

| **term** | **%coef +** | **%coef -** | **pooled pm** | **pooled std.error** |
|---|---|---|---|---|
| **(Intercept)** | 87 | 13 | 1.131 | 1.083 |
| **Body mass (g) ‡** | 67 | 33 | 0.382 | 0.867 |
| **Generation length ‡** | 94 | 6 | 0.707 | 0.511 |
| **Home range size (ha) ‡** | 86 | 14 | 0.512 | 0.496 |
| **Group size ‡** | 94 | 6 | 0.903 | 0.620 |
| **Brain volume ($cm^3$) ‡** | 56 | 44 | 0.172 | 1.068 |
| **Omnivore** | 53 | 47 | 0.027 | 0.362 |
| **Social system: Pair living** | 20 | 80 | -0.439 | 0.554 |
| **Social system: Polygynandry** | 59 | 41 | 0.159 | 0.725 |
| **Social system: Polygyny** | 65 | 35 | 0.288 | 0.789 |
| **Lifestyle: Arboreal** | 90 | 10 | 0.957 | 0.735 |
| **Lifestyle: Both** | 90 | 10 | 0.977 | 0.752 |
| **Insularity** | 92 | 8 | 1.147 | 0.928 |
| **Nocturnal** | 51 | 49 | 0.017 | 0.604 |

Table S20: Predictors of threat type 7: natural system modifications (binary) for 404 species with known threats using imputed traits values from Rphylopars (Goolsby et al., 2017). Posterior means and standard error pooled from 100 MCMCglmm models (one per imputed dataset and sampled phylogeny) with threshold error structures. Percentage of positive and negative coefficients come from 1,001 iterations retained across each of the 100 models (total retained iterations = 100,100). Each model included 10 traits as fixed effects and continuous variables ‡ were ln-transformed, centered with respect to the mean, and scaled by 2 standard deviations.

| **term** | **%coef +** | **%coef -** | **pooled pm** | **pooled std.error** |
|---|---|---|---|---|
| **(Intercept)** | 23 | 77 | -0.715 | 1.001 |
| **Body mass (g) ‡** | 88 | 12 | 0.881 | 0.750 |
| **Generation length ‡** | 34 | 66 | -0.146 | 0.375 |
| **Home range size (ha) ‡** | 57 | 43 | 0.056 | 0.323 |
| **Group size ‡** | 55 | 45 | 0.041 | 0.364 |
| **Brain volume ($cm^3$) ‡** | 36 | 64 | -0.353 | 0.997 |
| **Omnivore** | 13 | 87 | -0.297 | 0.270 |
| **Social system: Pair living** | 25 | 75 | -0.284 | 0.445 |
| **Social system: Polygynandry** | 15 | 85 | -0.546 | 0.538 |
| **Social system: Polygyny** | 6 | 94 | -0.846 | 0.558 |
| **Lifestyle: Arboreal** | 93 | 7 | 0.950 | 0.705 |
| **Lifestyle: Both** | 95 | 5 | 1.048 | 0.695 |
| **Insularity** | *0* | *100* | *-1.280* | *0.524* |
| **Nocturnal** | 15 | 85 | -0.490 | 0.496 |

**REFERENCES:**

DeCasien, A. R., Williams, S. A., & Higham, J. P. (2017). Primate brain size is predicted by diet but not sociality. *Nature Ecology & Evolution*, 1, 1-7.

Estrada, A., Garber, P. A., Rylands, A. B., Roos, C., Fernandez-Duque, E., Di Fiore, A., ... & Li, B. (2017). Impending extinction crisis of the world's primates: Why primates matter. *Science Advances*, 3, e1600946.

Galán-Acedo, C., Arroyo-Rodríguez, V., Andresen, E., & Arasa-Gisbert, R. (2019). Ecological traits of the world's primates. *Scientific Data*, 6, 1-5.

Goolsby, E. W., Bruggeman, J., & Ané, C. (2017). Rphylopars: fast multivariate phylogenetic comparative methods for missing data and within-species variation. *Methods in Ecology and Evolution*, 8, 22-27.

Harcourt, A. H., & Parks, S. A. (2003). Threatened primates experience high human densities: adding an index of threat to the IUCN Red List criteria. *Biological Conservation*, 109, 137-149.

IUCN. (2021). The IUCN Red List of Threatened Species Version 2021-1. IUCN Red List of Threatened Species. Retrieved from: https://www.iucnredlist.org/en.

Rowe, N., & Myers, M. (2011). *All the world's primates*. Primate Conservation, Inc. Retrieved from: https://www.alltheworldsprimates.org.
